# Body-wide genetic deficiency of poly(ADP-ribose) polymerase 14 sensitizes mice to colitis

**DOI:** 10.1101/2024.01.12.575356

**Authors:** Madhukar Vedantham, Lauri Polari, Anbu Poosakkannu, Rita G. Pinto, Moona Sakari, Jukka Laine, Petra Sipilä, Jorma Määttä, Heidi Gerke, Tiia Rissanen, Pia Rantakari, Diana M. Toivola, Arto T. Pulliainen

## Abstract

Inflammatory bowel disease (IBD) is a debilitating and relapsing chronic disease of the gastrointestinal tract affecting millions of people. Here, we investigated the expression and functions of poly (ADP-ribose) polymerase 14 (Parp14), an important regulatory protein in immune cells, using a biobank IBD patient cohort as well as two mouse models of colitis, i.e., the IBD-mimicking oral dextran sulfate sodium (DSS) exposure model, and the oral *Salmonella* exposure model. Parp14 was expressed in the human colon, by cells in the lamina propria, but, in particular, by the epithelial cells with a typical granular staining pattern in the cytosol. The same Parp14 staining pattern was evidenced in both colitis models. Body-wide genetic deficiency of Parp14 in C57BL/6N background sensitized mice to DSS colitis. The Parp14-deficient mice displayed increased rectal bleeding as well as stronger epithelial erosion, Goblet cell loss and immune cell infiltration. The absence of Parp14 did not affect the mouse colon bacterial microbiota based on PacBio long read sequencing. Also, the colon leukocyte populations of Parp14-deficient mice were normal based on flow cytometry. In contrast, we witnessed an altered transcriptional signature in Parp14-deficient mice with bulk tissue RNA-Seq. Gene Ontology (GO)-based classification of differentially expressed genes demonstrated that the colon transcriptional signature of Parp14-deficient mice was dominated by abnormalities in inflammation and infection responses both prior and after the 1-week DSS exposure. Overall, the data indicate that Parp14 has an important role in the maintenance of colon epithelial barrier integrity. The prognostic and predictive biomarker potential of Parp14 in IBD merits further investigation.

## INTRODUCTION

Inflammatory bowel disease [IBD, ulcerative colitis (UC) and Crohn’s disease (CD)] is a debilitating and relapsing chronic disease of the gastrointestinal tract affecting millions of people, mostly diagnosed in adolescence or early adulthood ^1^. The etiology of IBD remains unknown, but it appears to involve a genetic predisposition. Approximately 200 risk loci have been identified with major part in genes regulating immunological pathways controlling microbial recognition and killing, e.g., in *Nod2* gene ^2,3^. Indeed, infections with enteric pathogens such as *Salmonella* leading to gut microbiota dysbiosis and alterations of the intestinal immune responses have been debated as environmental factors ^2^. The disease culminates in dysfunction of the epithelial barrier, increase in immune cell infiltration, and elevated concentrations of the inflammatory cytokines ^1^. IBD can be controlled in only a subset of patients, e.g., with anti-TNF-α antibodies or leukocyte migration blocking anti-α4β7 integrin antibodies ^3^, highlighting the need to identify new drug targets. Patients that do not respond develop adverse effects, most notably increased risk of infections and colorectal cancer, thus requiring continuous medical monitoring, and sometimes removal of the affected tissues by surgery is required ^1,3^. There is a widespread consensus that a more individualised approach is required, with a need for more accurate prognostic and predictive biomarkers allowing more efficient patient stratification.

Poly (ADP-ribose) polymerase 14 (Parp14) is an enzyme of the Parp protein family ^4^. Parp family members catalyze ADP-ribosylation of macromolecules, i.e., proteins and nucleic acids, which refers to the covalent conjugation of an ADP-ribose moiety from nicotinamide adenine dinucleotide (NAD+) onto the substrate with simultaneous release of nicotinamide ^4^. Parp functions that are independent of the ADP-ribosyltransferase (ART) activity such as scaffolding and modification reversal are also important, as exemplified by the recent work on ADP-ribose binding and ADP-ribose hydrolysis activities of Parp14 macrodomains ^5,6^. Parp14 was identified as a Stat6 interacting protein ^7^, and accordingly acts as a transcriptional co-factor in interleukin-4 (IL-4)-induced Stat6-dependent gene expression in B cells ^7–10^. Parp14 has functions also in other lymphocytes, e.g., T cells from Parp14-deficient mice have difficulties to differentiate into IL-4/Stat6-dependent Th2 direction ^11,12^. Interestingly, Parp14 has negative downstream effects on interferon-γ (IFN-γ)- and Stat1-dependent cellular responses. Most notably, genetic Parp14 inactivation in macrophages skews them toward a pro-inflammatory IFN-γ-driven M1 phenotype while reducing the IL-4-driven M2 phenotype ^13^. Based on these opposing effects on IFN-γ- and IL-4-mediated gene expression, Parp14 has been debated to play a role in macrophage polarization ^13–15^. ADP-ribosylation of Stat proteins is one possible mechanism how Parp14 may regulate macrophage functions ^13,16^. Overall, Parp14 appears as an important regulator of a number of different immune cells functions.

We hypothesized that Parp14 is involved in the colon immune responses, and, if so, its malfunction could play a role in the onset or development of IBD. We explored the effect of body-wide genetic Parp14 deficiency in the robust murine model of IBD, i.e., in the 1-week oral dextran sulfate sodium (DSS) exposure colitis model ^17,18^. We also conducted a histological survey of Parp14 expression in the mouse colon in the 1-week oral DSS exposure colitis model, and, in parallel, in the single dose oral *Salmonella* exposure colitis model ^19^, and in the human colon using colon biopsies of IBD patients. The data highlight Parp14 as a protein highly expressed by epithelial cells, and having an important role in the maintenance of colon epithelial barrier integrity.

## RESULTS

### Parp14 is expressed by human epithelial cells

To examine Parp14 expression and localization in the human colon, we analyzed biobank-archived formalin-fixed paraffin-embedded (FFPE) endoscopic biopsies (Suppl. data file 1). The patients had histologically been diagnosed either with ulcerative colitis (UC, n=12) or Crohn’s disease (CD, n=7). In addition, we analyzed colon biopsies of patients with colonoscopy-justifying symptoms (n=9), but having a normal colon status upon histological scoring by a pathologist. Some cells in the lamina propria were positive for Parp14 (Fig. 1A-C), as judged by the staining with a commercial mouse monoclonal anti-Parp14 antibody (Fig. S1). However, the staining was most evident in the cryptal epithelial cells and in the epithelial cells facing the lumen of the colon (hereafter surface epithelial cells). We also observed that Parp14 was mostly localized in the cytoplasm and staining was granular both in the surface and in the cryptal epithelial cells (Fig. 1B, Fig. S2). There were some crypts, in particular in UC patients, at the immune cell infiltrated sites with thin mucous lining, lacking Goblet cells, which contained epithelial cells highly positive for Parp14 (Fig 1C). The Parp14 staining intensity was visually scored for two categories – surface epithelial cells and cryptal epithelial cells (0, faint or negligible staining to 3, high intensity). The biopsy sections showed stronger Parp14 staining in the surface epithelial cells when compared to cryptal epithelial cells in all patient groups (Fig. 1D). There was a trend towards less pronounced Parp14 staining in the surface epithelial cells of CD patients as compared to normal patients (*p*-value 0.0851). However, statistically significant differences (*p*-value <0.05) of Parp14 staining intensity between normal, UC and CD patients were not observed (Fig. 1D). In the context of cell cultures, we witnessed that Parp14 expression was at a negligible basal level in HeLa229 cells, i.e., in a human epithelial cell culture model (Fig. S3). However, upon incubation with some common inflammatory stimuli interferons IFN-a or IFN-g more Parp14 was detected in a time-dependent manner. Similar effects were also evident with THP-1 cells, i.e., in a human macrophage cell culture model (Fig. S3). Taken together, Parp14 is expressed in the human colon, in particular by the epithelial cells, with a characteristic granular staining pattern in the cytosol.

**Figure 1.**
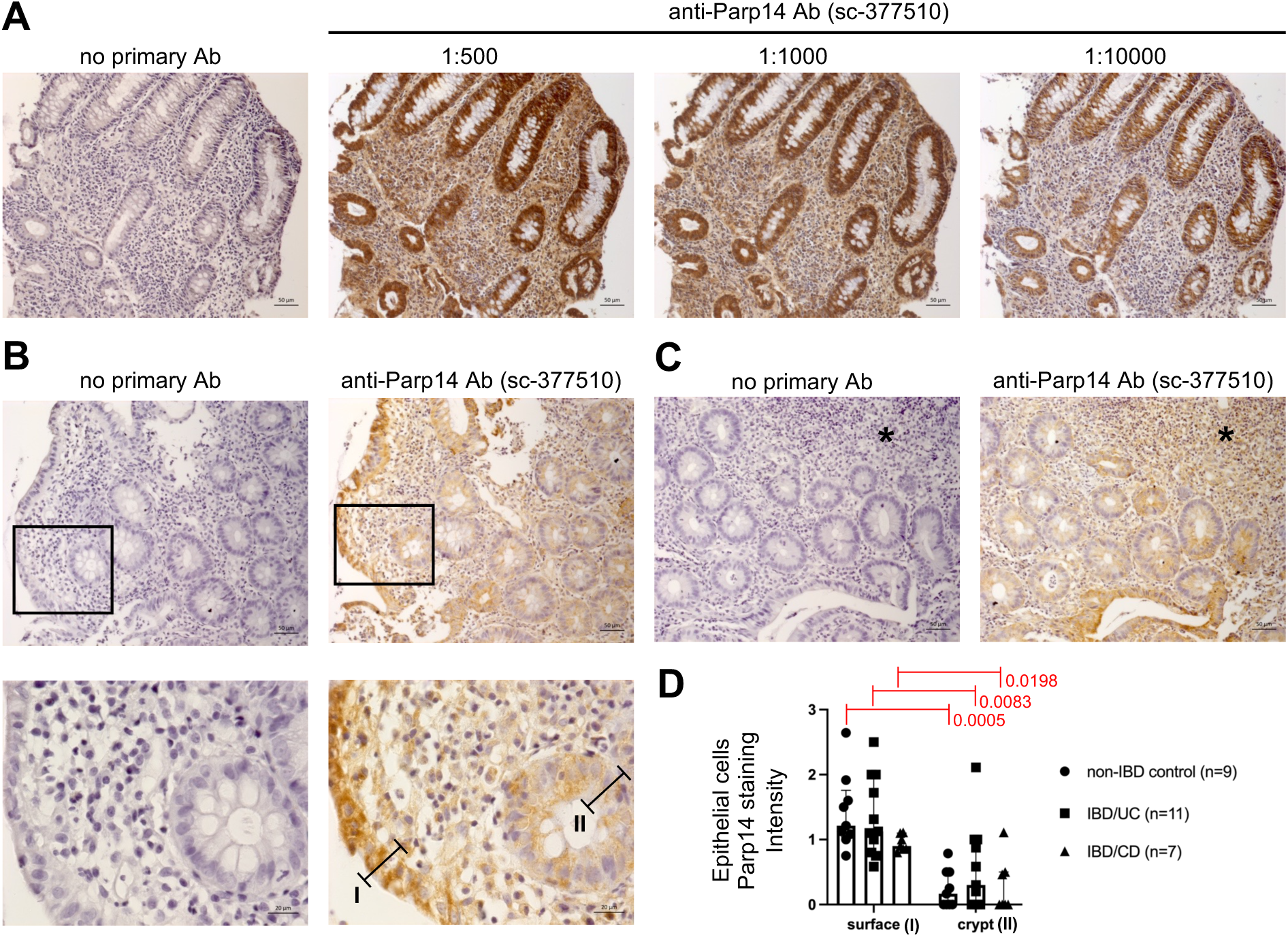
Parp14 expression and localization in the human colon. **A)** Immunohistochemistry (IHC) staining of Parp14 in an FFPE endoscopic biopsy section of an UC patient (TCML_VM_07A, descending colon, Suppl. data file 1). 20x objective images are shown. **B)** IHC staining of Parp14 in an FFPE endoscopic biopsy section of an UC patient (TCML_VM_21A, ascending colon, see Suppl. data file 1). 20x and 63x objective images are shown (1:10000 dilution of anti-Parp14 antibody). The bars 1 and 2 refer to the areas that were visually scored for Parp14 staining intensity (see Fig. 1D), i.e., I – surface epithelial cells facing the lumen of the colon, and II – cryptal epithelial cells. **C)** IHC staining of Parp14 in an FFPE endoscopic biopsy section of an UC patient (TCML_VM_21A, ascending colon, see Suppl. data file 1). 20x objective images are shown (1:10000 dilution of anti- Parp14 antibody). The star refers to a highly immune cell infiltrated site of the colon. **D)** Statistical comparison of Parp14 staining intensity in different patient groups. The analysis is based on staining intensity (10x objective images, 1:10000 dilution of anti-Parp14 antibody) in surface and cryptal epithelial cells that were scored as 0, faint or negligible staining to 3, high intensity. Statistical analysis was done with Kruskal-Wallis test.

### Parp14 is expressed by mouse colon epithelial cells

To examine Parp14 expression and localization in the mouse colon, we first analyzed FFPE distal colon sections derived from the DSS model executed in two genetic backgrounds - FVB/n and C57BL/6N. In the water control FVB/n mice, Parp14 staining was faint, whereas in the DSS-treated FVB/n mice (7 days, 2.5 % DSS in drinking water), the colon crypt epithelial cells were strongly Parp14 positive (Fig. 2A, Fig. S4). The staining was mostly granular in the cytoplasm similar to the human colon (see Fig. S2). The DSS-increased Parp14 staining was also evident, but not so clear, in the C57BL/6N background (Fig. 2A). Other cell types in the mouse colon also express Parp14. Some Parp14 positive cells in the colon wall of DSS-treated C57BL/6N mice were positive for the macrophage marker F4/80 (Fig. S5). Next, we analyzed FFPE proximal colon sections derived from the salmonellosis C57BL/6N model. In the PBS control C57BL/6N mice, Parp14 staining was faint, whereas in the *Salmonella*-infected mice (5 days post-infection), the colon crypt epithelial cells were strongly Parp14 positive (Fig. 2B). The staining was granular in the cytoplasm. Parp14 expression was also quantified at the transcriptional level. The level of Parp14 mRNA did not differ in the distal colon samples of water and DSS-treated FVB/n mice (Fig. 2C). However, Parp14 mRNA level was higher in the distal colon samples of DSS-treated C57BL/6N mice as compared to water controls (Fig. 2C). Moreover, Parp14 expression in the DSS treated C57BL/6N mice was higher in the distal colon as compared to proximal colon (Fig. 2C, *p*-value 0.0018). *Salmonella* infection (1 day and 5 days post-infection) also resulted in higher Parp14 mRNA levels as compared to PBS control in the proximal colon and cecum of C57BL/6N mice (Fig. 2D), but these differences were not statistically significant (lowest *p*-value 0.084, cecum PBS vs. day 5 infection). Taken together, Parp14 is expressed in the proximal and distal mouse colon, in particular by the epithelial cells, based on experimentation in two colitis models.

**Figure 2.**
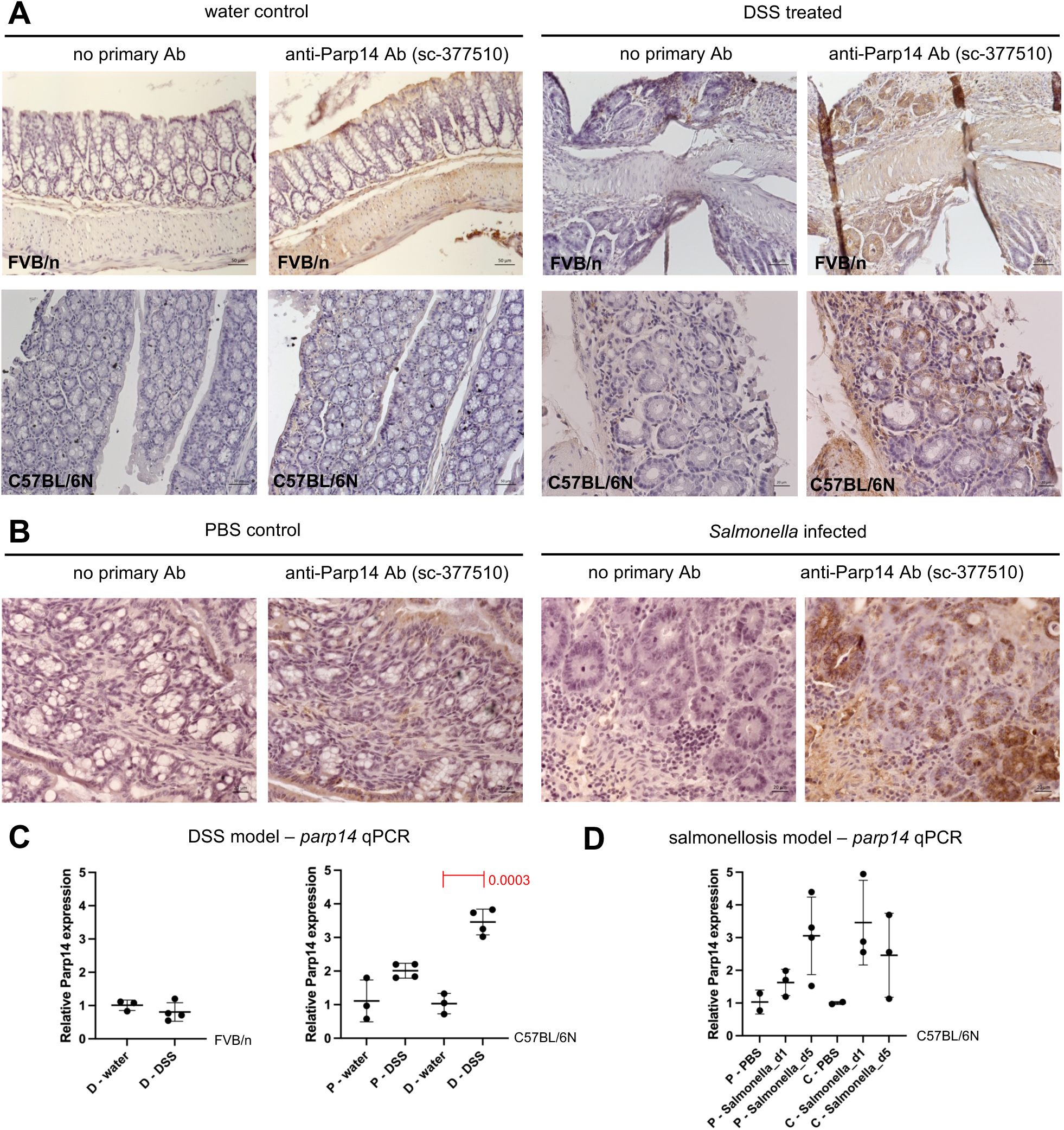
Parp14 expression and localization in the mouse colon. Parp14 expression was analyzed in two intestinal inflammation models – DSS and *Salmonella* colitis. **A**) DSS colitis IHC data. Parp14 IHC of distal colon of water control and DSS (2.5%, 7 days) treated FVB/n and C57BL/6N mice. 20x objective images are shown (1:500 dilution of anti-Parp14 antibody). **B**) Salmonellosis IHC data. Parp14 IHC of proximal colon of PBS control and *Salmonella* infected (5 days) C57BL/6N mice. 40x objective images are shown (1:1000 dilution of anti-Parp14 antibody). **C)** DSS colitis (day 8 of DSS treatment, see Fig. 3A) and **D)** salmonellosis (1- and 5-day post-infection) qPCR data on Parp14 expression (means with standard deviation, statistics with unpaired t-test). P – proximal colon; D – distal colon; C – cecum.

### Increased sensitivity of Parp14-deficient mice to DSS colitis

To analyze the physiological functions of Parp14 in colitis, we used the body-wide Parp14-deficient mice ^20^ backcrossed in-house to C57BL/6N background (Fig. S6). Littermates of wt and Parp14-deficient mice were treated with DSS and water as a control as described in Fig. 3A. We monitored the mice daily by quantifying their body weight, stool consistency and amount of blood in the stool. Oral administration of DSS caused loss in body weight in both wt and Parp14-deficient mice (Fig. 3B), in a statistically similar manner. The DSS-treated Parp14-deficient mice did not differ from wt mice in stool consistency (Fig. 3C). However, the Parp14-deficient mice suffered from increased rectal bleeding (Fig. 3D) in comparison to wt mice upon DSS treatment. This phenotype trend appeared at day 4, but it took until day 8 to reach statistical significance. The DSS treatment caused reduction of the colon length to a similar extent in wt and Parp14-deficient mice (Fig. 3E). We also performed a histopathological analysis of the distal and proximal regions of the colon. Hematoxylin and eosin (H&E)-stained FFPE sections were scored for edema, epithelial erosion, Goblet cell loss and immune cell infiltration. The absence of Parp14 did not cause apparent tissue damage in the water control mice, but it worsened the overall tissue damage observed upon the DSS treatment (Fig. 3F). Scoring of the different pathological variables revealed significantly stronger erosion and more pronounced immune cell infiltration in the distal colon of Parp14-deficient mice as compared to wt mice (Fig. 3G). Also, the Parp14-deficient mice suffered from more pronounced Goblet cell loss both in the proximal and distal colon (Fig. 3G). Taken together, the body-wide genetic deficiency of Parp14 sensitized mice to DSS colitis.

**Figure 3.**
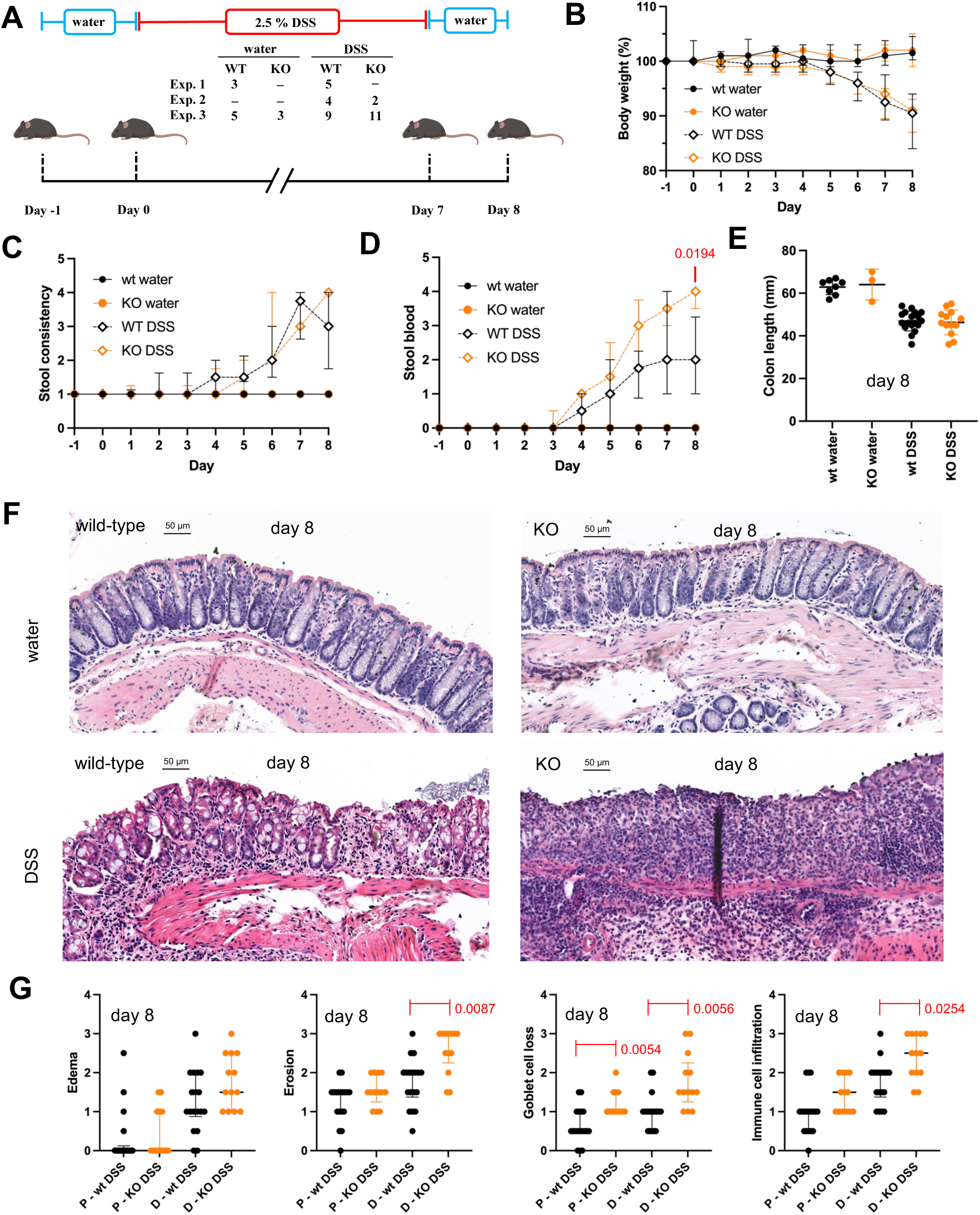
Body-wide genetic deficiency of Parp14 sensitizes mice to DSS colitis. **A)** Schematic description of the mouse experimentation. Animals from three different experiments were pooled for the analyses. **B)** Weight change of the mice during the course of the experiment relative to day -1 (medians with interquartile range). No statistical differences between wt and Parp14-deficient mice were detected. **C)** Stool consistency scoring (0 - normal feces to 4 - loose stool/diarrhea, medians with interquartile range). No statistical differences between wt and Parp14-deficient mice were detected. **D)** Stool scoring for blood (0 - no blood to 4 - bloody diarrhea with rectal bleeding, medians with interquartile range). Statistically significant difference between wt and Parp14-deficient mice is indicated in the figure. **E)** Colon length of the sacrificed mice (means with standard deviation). No statistical differences between wt and Parp14-deficient mice were detected. **F)** Images of the hematoxylin and eosin (H&E)-stained FFPE sections of distal colon samples of water and DSS treated wt and Parp14-deficient mice. 20x objective images are shown. **G)** Quantitation of pathological variables, i.e., tissue edema, epithelial erosion, Goblet cell loss and immune cell infiltration in proximal and distal colon (medians with interquartile range). Statistical differences between wt and Parp14-deficient mice are indicated in the figure. All the statistical analyses are described in the methods section.

### Absence of Parp14 does not affect the fecal bacterial microbiota

Bacterial colon dysbiosis in the Parp14-deficient mice could contribute to their increased sensitivity to DSS colitis. To explore this possibility, we used PacBio long-read sequencing of the nearly entire 16S rRNA gene and characterized the fecal bacterial microbiota during routine in-house colony maintenance. A total of nine bacterial phyla were found in the fecal pellets of wt and Parp14-deficient mice. The *Duncaniella*, *Faecalibaculum* and *Akkermansia* were the most dominant bacterial genera found with both mouse genotypes (Fig. 4A). To gain deeper insights, we first compared the operational taxonomic unit (OTU) richness (Chao1 index) and Shannon diversity index of the bacterial communities. There were no significant differences observed in Chao1 and Shannon alpha diversity indices between wt and Parp14-deficient mice (Fig. 4B), i.e., the genera richness did not differ. Secondly, to study if the genera composition differed between wt and Parp14-deficient mice, we used Principal Coordinate Analysis (PCoA) (Fig. 4C) as well as permutational analysis of variance (PERMANOVA) and beta dispersion analysis with Bray-Curtis dissimilarity metrics (data not shown). These beta diversity analyses demonstrated that the bacterial community composition of wt and Parp14-deficient mice did not statistically differ. Taken together, the absence of Parp14 does not appear to cause colon bacterial dysbiosis in Parp14-deficient mice during the routine in-house colony maintenance.

**Figure 4.**
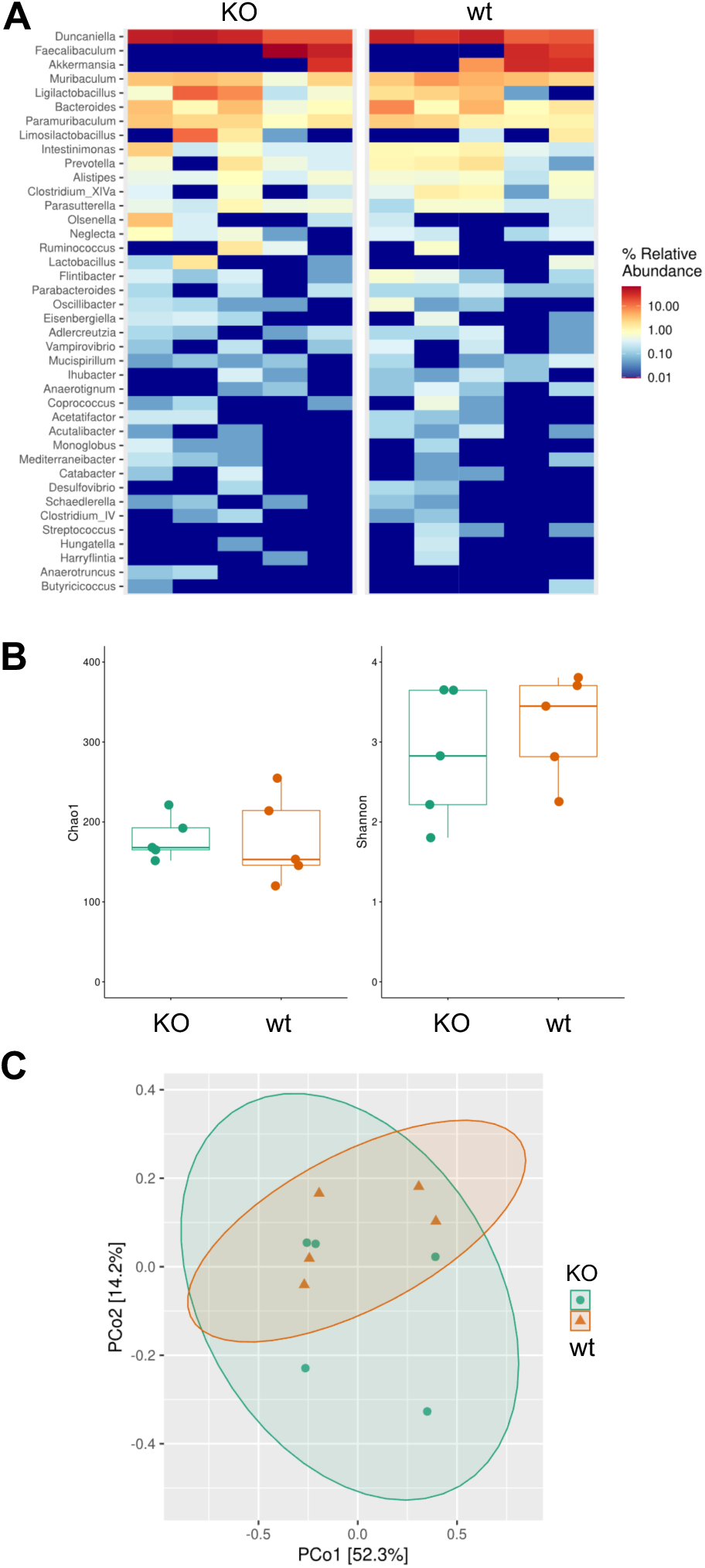
Body-wide genetic deficiency of Parp14 does not affect the fecal bacterial microbiota. **A)** A heat map showing the fecal bacterial microbiota profile of wt and Parp14-deficient mice. A total of 40 most abundant operational taxonomic units (OTUs), classified at the genus level, are listed in top to bottom with decreasing order of % relative abundance (OTU specific reads / all reads in a sample x 100). The OTUs are clustered at 97% similarity level. Each column represents a single individual mouse sample. **B)** The Chao1 and Shannon alpha diversity indices of the fecal bacterial microbiota richness differences in wt and Parp14-deficient mice. The OTUs are clustered at 97% similarity level. Each point represents an individual mouse fecal sample. No statistical differences between wt and Parp14-deficient mice were detected. **D)** The beta diversity output principal coordinate analysis (PCoA) of the fecal bacterial microbiota composition differences in wt and Parp14-deficient mice. No statistical differences between wt and Parp14-deficient mice were detected.

### DSS-treated Parp14-deficient mice have normal colon leukocyte populations

Abnormal colon immune cell population structure in Parp14-deficient mice could contribute to their increased sensitivity to DSS colitis. We used flow cytometry to detect different leukocytes from mid-colon, blood and spleen post-DSS and control treatments (Fig. 5, Fig. S7). We stained single-cell suspensions for cells of the myeloid lineage, i.e., granulocytes (neutrophils and eosinophils), monocytes, macrophages, dendritic cells, and cells of the lymphoid lineage (B and T cells). We did not detect significant differences in any of the leukocyte populations between water-treated wt and Parp14-deficient mice. The data implies that Parp14 is not a substantial driver of leukocyte development/homeostasis in mice (Fig. 5, Fig. S7). In both genetic backgrounds, and to similar extent, the DSS treatment increased the proportion of neutrophils and classical Ly6C^Hi^ and non-classical Ly6C^Low^ monocytes in the mid-colon compared to water-treated controls (Fig. 5). In a similar manner, DSS treatment significantly decreased the proportion of CD11c dendritic cells and the proportion of F4/80 macrophages (Fig. 5). Some genotype differences were evident in the blood and spleen. In the blood of DSS-treated Parp14-deficient mice, the proportion of circulating classical Ly6C^Hi^ monocytes was decreased as compared to wt mice (*p*-value 0.0322) (Fig. S7). Also, DSS did not induce a change in proportion of circulatory CD8 T cells in the Parp14-deficient mice (*p*-value 0.5457), although this took place in the wt mice (*p*-value of 0.0430) (Fig. S7). Furthermore, the absence of Parp14 caused a DSS-associated increase in the proportion of circulatory CD4 T cells (*p*-value 0.0242), which was not evidenced in the wt mice (*p*-value 0.6232) (Fig. S7). In the spleen, lower proportion of CD8 T cells (*p*-value 0.0205) was detected in the DSS-treated Parp14-deficient mice as compared to water control, while in the wt mice, this effect was not detected (*p*-value 0.3644) (Fig. S7). Overall, the data indicate that the DSS-treated Parp14-deficient mice had abnormal numbers of monocytes and T cells in the blood and spleen. The colon leukocyte populations of Parp14-deficient mice appeared normal.

**Figure 5.**
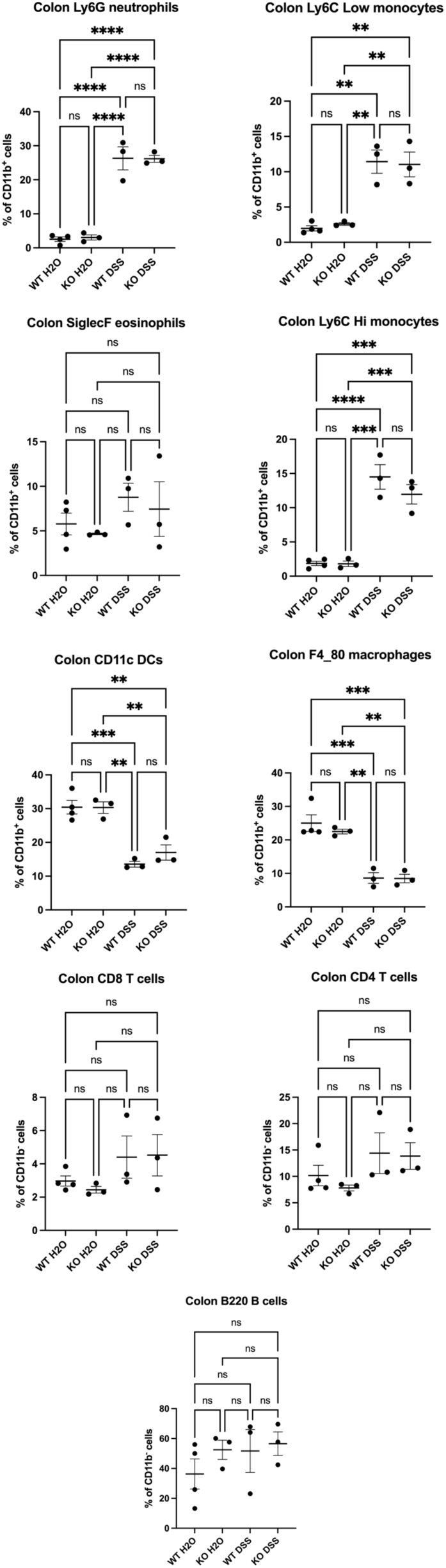
Body-wide genetic deficiency of Parp14 does not affect the colon leukocyte populations. Mid-colon single-cell suspensions post-DSS and control treatment at day 8 were stained for myeloid-derived cells, i.e., granulocytes (neutrophils and eosinophils), monocytes, macrophages, dendritic cells, and lymphoid-derived cells (B and T cells). Refer to the gating strategy in Fig. S7. Each dot represents one mouse. Data are presented as mean ±SEM. Statistical analysis is done with one-way ANOVA with the Bonferroni post-hoc test.

### Colon transcriptional signature of Parp14-deficient mice is skewed towards inflammatory signaling pathways

We performed bulk tissue RNA-Seq analysis of the mouse colon distal sections post-DSS treatment at day 8. The key data reliability metrics are shown in Fig. S8A-B. The heatmap cluster analysis indicated that there was differential gene expression between the compare groups (Fig. 6A). Numbers of genes detected to be expressed, as well as their differential expression metrics are shown in Fig. S8C-D, and Suppl. data file 2, respectively. First, we analyzed differentially expressed gene (DEG) differences in the water treated Parp14-deficient and wt mice. Based on the DEGs, we performed a Gene Ontology (GO) enrichment analysis. Only those DEGs that passed stringent filtering criteria were used in the GO enrichment analysis, i.e., up-regulated genes, log2(FoldChange) > 1 & padj < 0.05; down-regulated genes, log2(FoldChange) < -1 & padj < 0.05. Based on the genes up-regulated in Parp14-deficient mice (n=106), we did not identify any GO terms. In contrast, based on the genes down-regulated in Parp14-deficient mice (n=78), we identified 12 GO terms, which were all related to inflammation or infection (Fig. 6B-C). Out of all the 78 down-regulated genes, 34 (44 %) were mapped to these 12 GO terms (Fig. 6D). Specific examples included the C2 and C3 complement components, toll-like receptor 13, interleukin 7 receptor, chemokine (C-X-C motif) ligand 13, and vascular cell adhesion molecule 1. Next, we analyzed how wt and Parp14-deficient mice responded to DSS exposure. Of note, Parp14 was up-regulated in the wt mice post-DSS treatment at day 8 (1,606267 log2FoldChange, 3.044629-fold up in linear scale, padj 0,003323). Considerable differences between the DSS exposure-associated GO terms were detected between wt and Parp14-deficient mice (Fig. 6E-F, Suppl. data file 3). Firstly, the wt mice were more dynamic in respect of the numbers of DEG GO terms (Fig. 6E). Secondly, the up-regulated transcriptional response of Parp14-deficient mice was dominated by GO terms, which were related to inflammation or infection (Fig. 6F). Overall, the data imply that colon transcriptional signature of Parp14-deficient mice was skewed towards inflammatory signaling pathways both prior and after the DSS exposure.

**Figure 6.**
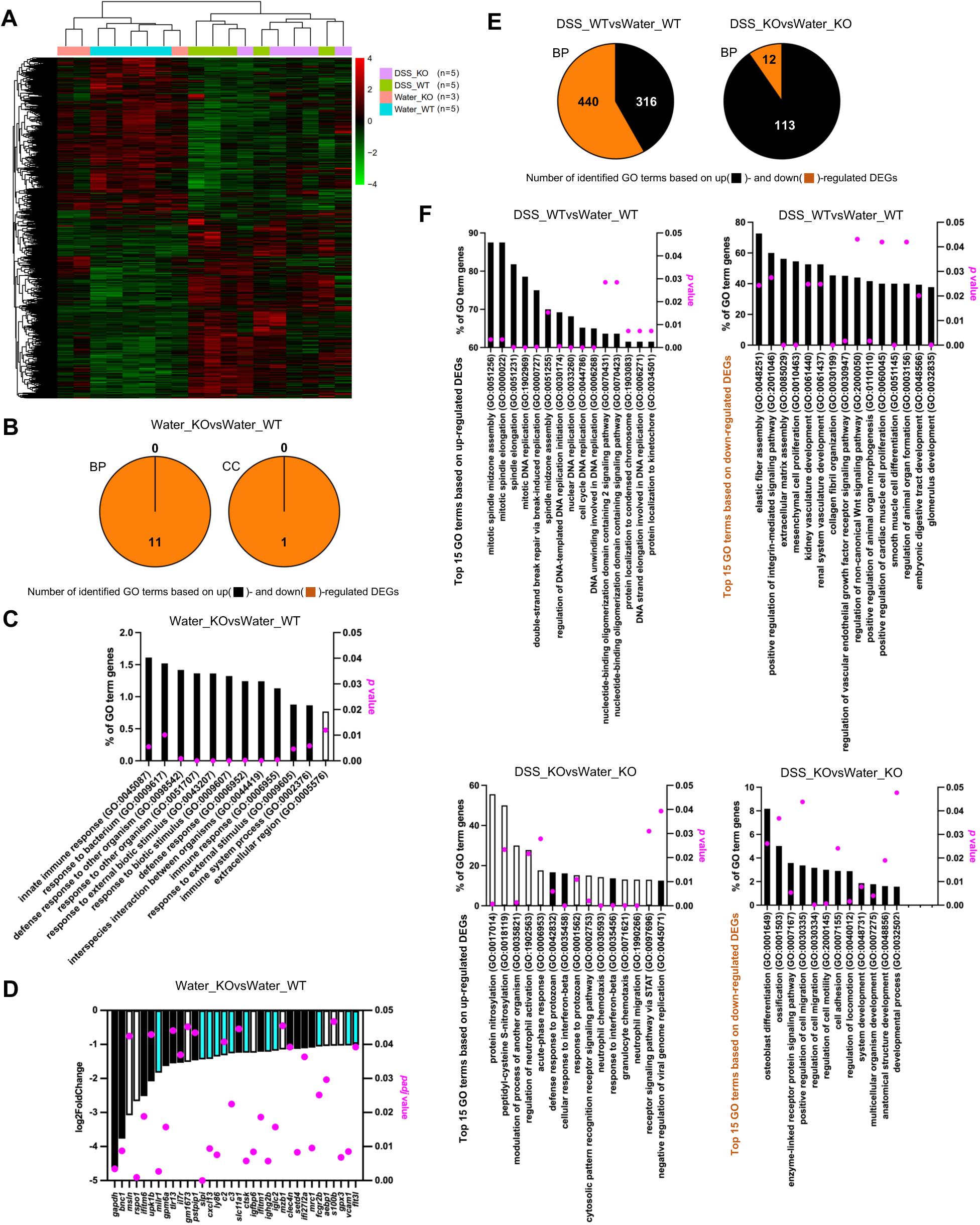
Colon transcriptional signature of Parp14-deficient mice is skewed towards inflammatory signaling pathways. **A)** Heatmap cluster analysis for differential gene expression between the samples. Red color indicates genes with high expression levels, and green color indicates genes with low expression levels. **B-C)** Gene Ontology (GO) enrichment analysis with genes that were down-regulated in the water treated Parp14-deficient mice as compared to the water treated wt mice. Based on the DEGs (Suppl. data file 2), GO terms were searched with binomial test using the Bonferroni correction. The bars refer to % of the detected genes out of all the genes in a given GO term. The black bars refer to the BP (Biological Process) and the white bar to CC (Cellular Component) GO terms. **D)** The down-regulated genes leading to the identification of the 12 GO terms (see Fig. 6B-C). Out of all the 78 down-regulated genes (Water_KOvsWater_WT), 34 (44 %) were mapped to these 12 GO terms. The white bars refer to the DEGs leading uniquely to this one CC GO term identification. The cyan bars refer to the DEGs leading both to this one CC and 11 BP GO term identifications. The black bars refer to the DEGs leading uniquely to the 11 BP GO term identification. **E-F)** Gene Ontology (GO) enrichment analysis with genes that were differentially expressed in the DSS treated Parp14-deficient and wt mice as compared to water treated controls. Based on the DEGs (Suppl. data file 2), GO BP terms were searched with binomial test using the Bonferroni correction. The bars refer to % of the detected genes out of all the genes in a given GO term. The white bars refer to the unique DSS response of the Parp14-deficient mice.

## DISCUSSION

Parp14 is an important regulator of immune cell functions, in particular, in lymphocytes and monocyte/macrophages ^7–16^. We explored the effect of body-wide genetic Parp14 deficiency in the murine model of IBD, i.e., in the 1-week oral DSS exposure colitis model ^17,18^. We also conducted a histological survey of Parp14 expression and localization in mouse colon, in the DSS colitis model, and, in parallel, in the oral *Salmonella* exposure colitis model ^19^, and in the human colon using biobank-archived colon biopsies of IBD patients.

The histological inspection of our IBD patient biopsies (UC, n=11; CD, n=7; control, n=9) demonstrated that Parp14 staining was most evident in the cryptal epithelial cells and in the epithelial cells facing the lumen of the colon, i.e., surface epithelial cells. We also observed that Parp14 was mostly localized in the cytoplasm and staining was granular. The granular staining pattern might be related to Parp14 localization into processing bodies (P-bodies), as reported with *in vitro* co-localization analysis of endogenous DEAD-box helicase 6 (DDX6) P-body marker and transfected GFP-Parp14 in HEK293T cells ^21^. Overall, more intense Parp14 staining was detected in the surface as compared to the cryptal epithelial cells in all three patient groups. This staining pattern could be related to the fact that the surface epithelial cells are more exposed to the colon luminal content, including the microbiota. Statistically significant differences were not detected in Parp14 staining intensity between the three patient groups. However, there were some crypts particularly in UC patients, at the immune cell infiltrated sites with thin mucous lining, lacking Goblet cells, which contained epithelial cells highly positive for Parp14. This could be related, in addition to luminal content assault, to stimulus from the lamina propria cells, e.g., the Parp14 up-regulating interferons ^15^ (see Fig. S3). While our study was in progress, a bulk colon RNA-Seq-driven study was published implying *parp14* as a potential driver gene in UC progression from limited disease to extended disease^22^. Histologically, the frequency of lamina propria cells with high Parp14 nuclear staining (rabbit polyclonal anti-Parp14, ab224352, epitope unknown), some of which were CD68 positive macrophages, was significantly higher in patients with UC that subsequently extended (n=6) as compared with non-extenders (n=5) ^22^. This nuclear localization of Parp14 most likely reflects its potent role as transcriptional regulator ^7,8,15^. We also detected Parp14 positive lamina propria cells but based on our cohort and used mouse monoclonal anti-Parp14 antibody (see Fig. S1), the staining intensity in the cytosol of epithelial cells dominated the overall histology. Of note, the histology images of Argmann and co-workers ^22^ also revealed granular cytosolic Parp14 staining in the human colon epithelial cells, but this finding was not addressed in their study. More recently, a bioinformatics-driven study on publicly available transcriptome data identified *parp14* as a ferroptosis-related gene upregulated in UC ^23^. We also found out highly significant up-regulation of *parp14* in two public repository UC and CD IBD cohort transcriptomes ^24,25^ (Fig. S9). Overall, Papr14 appears as a possible new mRNA or protein-based colon biopsy biomarker in IBD. However, more elaborate cohorts should be executed to evaluate its prognostic and predictive potential in IBD.

To analyze the physiological functions of Parp14 in colitis, i.e., in the 1-week DSS exposure colitis model, we used the body-wide Parp14-deficient mice ^20^ backcrossed in-house to C57BL/6N background. Oral administration of DSS to mice via drinking water induces severe colitis characterized by weight loss, bloody diarrhea, loss of epithelial cells and neutrophil infiltrations, resembling flares in human UC ^17,18^. DSS is believed to cause direct damage to the epithelial cells followed by strong immune system activation by dissemination of the proinflammatory colon luminal content such as bacteria into the sub-epithelial space ^17,18^. Most likely, the strong pro-inflammatory response including oxidative stress exacerbates the deleterious effects of DSS. Our key human histological findings were replicated in the DSS and also in *Salmonella* colitis models, in particular, the strong granular cytosolic Parp14 staining in the cryptal epithelial cells of an inflamed colon. Also, based on the DSS model, many of the Parp14 positive mouse lamina propria cells were F4/80 positive, and thereby classify as macrophages. As compared to wt mice, Parp14-deficient mice displayed increased rectal bleeding as well as stronger epithelial erosion, Goblet cell loss and immune cell infiltration, in particular, in the distal colon upon DSS exposure. Extreme epithelial erosion was frequently evidenced in the Parp14-deficient mice (see Fig. 3F). The body-wide genetic deficiency of Parp14 sensitized mice to DSS colitis.

Abnormal colon microbiota or abnormal colon immune cell population structure could explain the increased sensitivity of Parp14-deficient mice to DSS colitis. We did not detect differences in the fecal microbiota composition between the wt and Parp14-deficient mice during our routine colony breeding. Therefore, it appears unlikely that pro-inflammatory stimulus of a differential colon microbiota explains the observed phenotypes upon DSS exposure. Similarly, we did not detect differences in the colon monocyte/macrophage, dendritic cell, eosinophil, neutrophil, B cell and helper as well as killer T cell populations between the wt and Parp14-deficient mice after the DSS exposure or control water treatment. In respect of neutrophils, these data were somewhat unexpected, as more pronounced immune cell (neutrophil) infiltration was scored for Parp14-deficient mice under the DSS exposure with colon histological sections. This discrepancy could be explained by the mid vs distal locations of the analyzed samples. Unfortunately, the scarcity of sample material did not allow us to conduct flow cytometry-based leukocyte profiling of the distal colon. It is noteworthy that some leukocyte population structure differences between the wt and Parp14-deficient mice were detected in the peripheral tissues, namely in the spleen and blood monocyte/macrophage, T cell and B cell populations. In respect of lymphocytes, these data might reflect the early findings of T and B cell sub-population abnormalities in Parp14-deficient mice, e.g., lower marginal zone and higher follicular splenic B cell proportion as compared to the wt littermates ^20^. However, the detected statistically significant effects in our study were always to the same direction as evidenced in the other genotype (see Fig. S7). For example, in the blood of DSS-treated Parp14-deficient mice, the proportion of circulating classical Ly6C^Hi^ monocytes was decreased as compared to wt mice. Yet, similar kind of trend without statistical support was already apparent in the control water treated mice. We think that more experimentation with bigger numbers of mice would be needed to substantiate these findings. Moreover, it is the colon leukocyte population structure that intuitively should have the major impact, if any, on the observed phenotypes upon DSS exposure. Based on the data such differences do not exist between wt and Parp14-deficient mice, not upon the DSS exposure or control water treatment. The data implies that Parp14 is not a substantial driver of leukocyte development/homeostasis in mice, and that Parp14 does not affect the mouse colon leukocyte population structure upon DSS exposure.

Abnormal functions of the different colon cells prior the DSS exposure could explain the increased sensitivity of Parp14-deficient mice to DSS colitis. To experimentally address this possibility, we determined transcriptomes as the proxy of cellular functions using bulk tissue RNA-Seq. We clustered differentially expressed genes (DEGs) to cellular and physiological functions by Gene Ontology (GO) enrichment analysis, and compared the water treated Parp14-deficient mice to wt mice. This comparison tells us of the possible transcriptional priming of the colon wall of Parp14-deficient mice to DSS exposure. A relatively low number of DEGs that passed our filtering criteria were detected (n=184). Only the down-regulated genes (n=78) resulted in GO term identifications. Interestingly, all the identified 12 GO terms were related to inflammation or infection. Key examples of the down-regulated genes were the three interferon-inducible genes (*ifitm1*, *ifitm6*, *ifi27l2a*), two pattern recognition receptor genes (*tlr13*, *clec4n*), one chemokine gene (*cxcl13*), and one cytokine receptor gene (*il7r*). We also detected down-regulation of two complement genes (*c2*, *c3*) encoding for the C2 classical pathway component and for the central C3 activatory hub of all the complement pathways. At the same time one alternative pathway gene (*cfb*) encoding for factor B acting upstream of C3 was strongly up-regulated in the Parp14-deficient mice. Due to its central role in the regulation of inflammation, complement system has also gathered interest in the IBD research field ^26^. For example, the C3-deficient mice displays decreased intestinal inflammation in the acute ^27^, whereas increased intestinal inflammation in the chronic ^28^, DSS exposure colitis model. It is therefore tempting to speculate that the detected complement gene expression alterations in our mice prior to DSS exposure contribute to the observed phenotypes upon DSS exposure. It is noteworthy that we did not detect signs of elevated pro-inflammatory status of the colon wall in the Parp14-deficient mice. This indicates that the colon wall epithelial cell barrier was not significantly compromised in the Parp14-deficient mice prior the DSS exposure. Taken together, the colon transcriptional signature of Parp14-deficient mice was dominated by abnormalities in inflammation and infection response prior the DSS exposure.

The increased sensitivity of Parp14-deficient mice to DSS colitis could also be explained by functional abnormalities of the different colon cells during the DSS exposure. We therefore analyzed how wt and Parp14-deficient mice responded to the DSS exposure, in particular, what genes got up-regulated. Considerable differences between the DSS exposure-associated GO terms were detected. First, the up-regulated DSS response in wt mice was dominated by DNA replication- and cell division-related GO terms. This implies that the wt mice were more or less coping with the DSS exposure by enhancing the colon wall renewal. Inflammation- and infection-related GO terms were also identified, but in the GO term ranking they appeared low. In contrast, the up-regulated DSS response of Parp14-deficient mice was dominated by inflammation- and infection-related GO terms. All the top 15 scoring GO terms were related to inflammation and infection, and 11 of these were not detected at all in the wt mice. This implies, in accordance with our disease severity scores, that the Parp14-deficient mice were more or less beyond the epithelial renewal attempts while trying to merely cope with the strong sub-epithelial assault of the luminal content. However, it remains elusive if the detected transcriptomic profile at day 8 in the Parp14-deficient mice was driving the pathology or if it merely reflected it. Based on *in vitro* experimentation with macrophages, Parp14 deficiency induces the pro-inflammatory M1 polarization upon IFN-γ stimulation, whereas, at the same time, it suppresses the anti-inflammatory M2 polarization upon IL-4 stimulation ^13^. It is tempting to speculate that Parp14 deficiency in the macrophages had converted the colon wall into a more tissue damaging environment under the DSS exposure. However, our data demonstrated high Parp14 expression and thereby highly plausible functions also in the epithelial cells. As our bulk tissue RNA-Seq data lacks the cell identity information both spatially and temporally, single cell spatial transcriptomics could in future provide better understanding of the Parp14 functions. This information could also allow creation of hypotheses for feasible cell culture-based experiments to understand at molecular level how and in which cell type Parp14 executes its functions.

Parp14 is subject of an active drug development campaign. Compounds that target the NAD+ binding pocket with subsequent ART activity inhibition, e.g., ^14,29^, or that target the NAD+ binding pocket with subsequent Parp14 proteolysis (proteolysis targeting chimera) ^30^ have been reported. Also, one line of research has focused on compounds that target the Parp14 macrodomains, e.g., ^31–33^, involved in the binding of Parp14 to ADP-ribose conjugates and removal of them ^5,6^. Therefore, pharmaceutical perturbation of Parp14 functions is being explored at multiple levels from the overall Parp14 amount to its ART activity, and from Parp14-mediated ADP-ribose-dependent molecular scaffolding to reversal of ADP-ribosylation dependent cell signalling events. First of all, these studies are fuelled by the genetic findings where body-wide deficiency of Parp14 in mice diminished the intensity of allergic reactions ^11,12,34^. To this end, one ART activity inhibitor was recently shown to reduce pulmonary allergic response in mice ^35^ and one ART activity inhibitor entered clinical trials in atopic dermatitis (Phase I, NCT05215808). Secondly, *in vitro* genetic findings have demonstrated that Parp14 inactivation in macrophages skews them toward a pro-inflammatory IFN-γ-driven M1 phenotype while reducing the IL-4-driven M2 phenotype ^13^. Accordingly, one ART activity inhibitor has been explored to reverse the immunosuppressive M2 phenotype of tumor-associated macrophages^14^. Also, one recent publication reported on the potency of different Parp14 inhibitors to overcome the chronic IFN-γ-associated resistance to anti-PD-1 immune checkpoint inhibitor therapy in melanoma ^36^. As for now, no published data is available on Parp14 inhibitors in pre-clinical models of IBD. Our study with body-wide Parp14 knockout mice indicates that Parp14 inhibition could be detrimental in IBD. However, we argue that more elaborate experimentation is required, e.g., by using the chronic DSS exposure colitis model, and possibly other chemical models of IBD such has the oxazolone model ^17^. Indeed, our genetic experiments tell little of the therapeutic potential of small molecular weight compounds to reverse the symptoms of already established pathologies.

In summary, our study in the DSS colitis mouse model of IBD ^17,18^ provides compelling evidence that Parp14 has an important role in the maintenance of colon epithelial barrier integrity. The Parp14-deficient mice had increased sensitivity to DSS colitis. The strong colon epithelial cell expression pattern of Parp14 detected in our study in human and mice, and while our study was in progress by Argmann and co-workers in human ^22^, indicates that Parp14 may have functions also in non-immune cells, in addition to its important regulatory functions in immune cells ^7–16^. Our finding of the strong focal histological staining pattern of Parp14 in some IBD patients together with the recent mRNA level indications of Parp14 as a potential driver gene in UC ^22,23^ highlights the need for bigger cohorts to assess Parp14 as a new prognostic or predictive biomarker in IBD.

## ACKNOWLEDGEMENTS

This work was financially supported by the Research Council of Finland grants with project numbers 295296 and 329252 to ATP as well as 315139 and 332582 (including InFLAMES Flagship Programme, 337531 and 357911) to DMT and the 1-year Finnish Cultural Foundation personal grant 00231206 to MV. In addition, MV has received a 2-year salary package from the Turku Doctoral Programme of Molecular Medicine (TuDMM). Mika Savisalo, Merja Lakkisto, and personnel of the Histology core facility (Institute of Biomedicine, University of Turku, Turku, Finland) are acknowledged for technical support.

## MATERIALS AND METHODS

### Expression and purification of GST-Parp14 fragment

***i) Plasmid.*** Synthetic DNA fragment (Eurofins Genomics) encoding for amino acids 291-358 of the human Parp14 (Uniprot Q460N5), i.e., epitope of the sc-377150 antibody (Santa Cruz Biotechnology), was cloned with BamHI and XhoI into pGEX-6-P1 (N-terminal GST-tag, pGEX-6-P1-PARP14^291–358^). ***ii) Expression.*** The pGEX-6-P1-PARP14^291–358^ plasmid was transformed into BL21(DE3) (Novagen) and selected overnight at 37°C on Luria-Bertani (LB) agar with 200 µg/mL of ampicillin. Next day, one colony was inoculated to 5 mL of liquid LB with 200 µg/mL of ampicillin, and grown overnight at 37°C with shaking. Next day, the bacteria were sub-cultured to LB 1:100 with 200 µg/mL of ampicillin and grown at 37°C with shaking until the OD600 reached 0.5 upon which isopropyl β-D-1-thiogalactopyranoside (IPTG) was added to 500 µM. The culture was grown at room temperature with shaking for 4 hours. Bacteria were collected by centrifugation and were frozen to -80°C. ***iii) Purification.*** Lysozyme (Sigma, L6876) and protease inhibitors (Thermo Fisher Scientific, A32965) were added to 0.5 mg/mL and 1 tablet / 50 mL, respectively, in the thawed biomass in lysis buffer [50 mM Tris (pH 8.0), 150 mM NaCl, 2 mM dithiotreitol (DTT)]. The sample was sonicated and clarified by centrifugation. Supernatant was incubated with Protino Glutathione Agarose 4B beads (Macherey-Nagel, 745500) in a rotatory shaker overnight at +4°C. The beads were applied to an empty plastic chromatography column and washed with gravity flow using 50 mM Tris (pH 8.0), 150 mM NaCl, 2 mM DTT. The GST-PARP14^291–358^ protein was eluted with 50 mM Tris (pH 8.0), 150 mM NaCl, 2 mM DTT, 25 mM glutathione (Sigma, G4251), aliquoted and stored at -80°C.

### Patient material and Parp14 immunohistochemistry

FFPE biopsy specimens sectioned at 5 µm thickness from anonymous IBD (UC and CD) vs control patients were obtained from the Finnish biobank system (https://site.fingenious.fi/en/, Auria Biobank, Turku, Finland, project number ab20-3646). The FFPE sections were stained for Parp14 with mouse monoclonal antibody (sc377150, Santa Cruz Biotechnology, dilutions 1:500, 1:1000 and 1:10000) and detected using the mouse specific HRP-DAB (ABC) detection immunohistochemistry (IHC) kit (ab64259, Abcam). Tissue sections were air dried for 2 h at room temperature, placed in 37°C incubator overnight, deparaffinized in xylene and rehydrated with alcohol gradients. Endogenous peroxidase was blocked with ‘Peroxidase blocking solution’ provided with the IHC kit. The sections were immersed in prewarmed 10 mM Na-citrate buffer (pH 6.0) and kept in boiling water bath for 20 minutes. After antigen retrieval, sections were rinsed in PBST (PBS with 0.01% Tween-20) followed by BSA (5% w/v in PBST) blocking for 1 h at room temperature to reduce nonspecific binding of the antibodies. The primary anti-Parp14 antibody in BSA (5% w/v in PBST) was added to tissue sections and incubated overnight at 4°C in a humidified chamber. Post incubation with primary antibody, tissue sections were rinsed in PBST twice for 5min each and incubated with biotinylated anti-mouse secondary antibody (provided with IHC kit) for 1h at room temperature. Streptavidin-HRP conjugate was added and then stained using 3-3’ Diaminobenzidine (DAB) as a chromogen (both solutions were provided with IHC kit). Harris hematoxylin was used as a nuclear counterstain. Sections without incubation of primary antibody served as negative controls. Mounting was done using Histo-Clear and sections were allowed to sit at room temperature for 12h before imaging. Imaging was done using Zeiss AxioImager M1 microscope with 5x, 20x, 40x oil or 63x oil objective lenses. Two persons, one being a pathologist, independently and blind for sample group, scored the stained sections (1:10000 of anti-Parp14 antibody) by visual inspection using a bright field microscope. The Parp14 staining intensity (0 - none, 1 - faint and irregular, 2 - mild and regular, 3 - strong and highly regular) was scored for the surface epithelial cells and the cryptal epithelial cells (see Fig. 1B). The score of one particular patient means average of scores from all the available tissue sections across the gastrointestinal tract, which varied from patient to patient. The scores of the two persons were averaged. Differences between cryptal and surface epithelial cell staining intensities in patient groups were summarized with descriptive statistics and studied by Kruskal-Wallis test. The normality of variables was evaluated visually and tested with the Shapiro-Wilk test. Due to the non-normality of the continuous variables, nonparametric methods were used. The statistical significance level was set at 0.05 in all tests (two-tailed). The analyses were performed using the SAS system, version 9.4 for Windows (SAS Institute Inc., Cary, NC, USA).

### Validation of the anti-Parp14 antibody

***i) Western analysis.*** Synthetic DNA fragment (Eurofins Genomics) encoding for the entire mouse Parp14 (Uniprot Q2EMV9) was cloned into pcDNA3.1-Hygro(-)-based human EGFR-HA expression plasmid ^37^. The human EGFR insert was replaced with the mouse Parp14 insert before the HA-tag-encoding area using NheI and NotI (C-terminal HA-tag, pcDNA3.1-Hygro(-)-mParp14). The pcDNA3.1-Flag-Parp14 plasmid to express human Parp14 (N-terminal Flag-tag) has previously been described ^38^. HEK293T cells grown in DMEM (21969035, Thermo Fisher) + 10% FBS (S181B-500, Biowest) + 2mM L-glutamine (25030081, Gibco) + 25mM HEPES (15630-056, Gibco) were seeded in 10 mL volumes in 10 cm dishes (1×10^6^ cells / 10 cm dish) in the late afternoon. The next morning fresh media was exchanged and the cells were transfected with 4 µg of plasmid DNA using calcium-phosphate transfection reagents (prepared in-house). At 24 h post-transfection, cells were washed with ice-cold sterile PBS and lysed in modified RIPA buffer (50mM Tris pH 7.5, 400mM NaCl, 1% NP-40, 0.5% sodium deoxycholate, 0.1% sodium dodecyl sulfate, 1mM EDTA), 75 µM Tannic acid (PARG inhibitor, 403040, Sigma Aldrich), 40 µM PJ-34 (PARP inhibitor, J64413, Fischer Scientific), Pierce protease-phosphatase inhibitor (A32961, Thermo Fisher Scientific) and cleared by high-speed centrifugation at 4°C. Protein concentration was measured from the supernatants with Bradford protein assay. The heated (10 min, 95 °C) samples in Laemmli loading dye were run on SDS-PAGE (30 µg / lane) and transferred to nitrocellulose membrane, followed by blocking with 5% (w/v) skimmed milk in Tris-buffered saline with 0.1% Tween 20 detergent (TBST). Primary antibody solution was prepared with 5% (w/v) skimmed milk in TBST with anti-Parp14 mouse monoclonal antibody (1:500, sc377150, Santa Cruz Biotechnology) with enough volume for two membranes (epitope competition vs control). With the epitope competition, one-half of this primary antibody solution was incubated in a rotary platform for 30 min at room temperature with GST-Parp14^291–358^ protein (concentration 15x more than with the primary antibody) prior to incubation with the membrane in a rotary platform for 48 h at 4°C. Membranes were washed thrice with TBST containing 5% (w/v) skimmed milk for 10 min each time. Membranes were incubated in TBST containing 5% (w/v) skimmed milk with goat anti-mouse IgG conjugated to horseradish peroxidase (HRP) (1:2000, 1010-05, SouthernBiotech) for 3h at 4°C in a rotary platform and washed thrice with TBST for 10 min each time. Membranes were subsequently developed with WesternBrightECL (Advansta) and imaged on ImageQuant LAS 4000 (GE Healthcare). Post imaging, same membranes were probed with anti-beta actin HRP conjugate (sc-47778 HRP, Santa Cruz Biotechnology). Membranes were developed and imaged as above. ***ii) Immunohistochemistry.*** FFPE sections of an UC patient (TCML_VM_28A, ascending colon, see Suppl. data file 1) were prepared for IHC as described above. The primary anti-Parp14 antibody in BSA (5% w/v in PBST) was added to tissue sections and incubated overnight at 4°C in a humidified chamber. The epitope competition slide was processed in parallel in a similar manner, but the primary anti-Parp14 antibody solution was pre-incubated for 30 min 4°C using 15x more GST-Parp14^291–358^ protein than the primary antibody. Secondary antibody treatment and counterstaining were the same as described above. Imaging was done using Zeiss AxioImager M1 microscope with 5x, 20x, and 40x oil objective lenses.

### Western analysis of Parp14 expression in cell cultures upon cytokine stimulation

HeLa229 human cervical adenocarcinoma cells (CCL-2.1, ATCC) grown in DMEM (21969035, Thermo Fisher) + 10% FBS (S181B-500, Biowest) + 2mM L-glutamine (25030081, Gibco) + 25mM HEPES (15630-056, Gibco) were seeded in 3 mL volumes in 6-well plates (3×10^5^ cells / well). THP-1 human monocyte/macrophage cells (TIB20, ATCC) grown in RPMI 1640 (Lonza) + 10% FBS (S181B-500, Biowest) were differentiated using 10 ng/mL PMA (phorbol 12-myristate 13-acetate, P8139, Sigma) for 24 h. THP-1 cells were washed twice with sterile PBS and fresh RPMI media without PMA was used to seed cells (2×10^5^ cells/well) in 3mL volumes in 6-well plates. Next morning, cells (HeLa229 and THP-1) were washed again with PBS and exchanged with fresh RPMI media supplemented with 100 ng/mL of *E. coli* 0111: B4 strain derived LPS (tlrl-3pelps, InvivoGen), 100 U/mL of IFN-γ (PHC4031, Thermo Fisher Scientific), 100 U/mL of IFN-α (SRP4596, Sigma Aldrich) or 10 ng/mL of TNFα (210-TA, R&D systems). Wells with fresh media without any inflammatory stimuli served as control. At times 2h, 8h, 12h, 24h and 48h post-stimulation, cells were washed twice with sterile PBS, trypsinized and lysed using ice-cold NP-40 lysis buffer (50mM Tris pH 7.5, 1% NP-40, 1x Pierce protease-phosphatase inhibitors with EDTA). Sample preparation for SDS-PAGE and Western analysis was done as described above. After blocking with TBST containing 5% (w/v) skimmed milk, membranes were probed with anti-Parp14 mouse monoclonal antibody (1:500, sc-377150, Santa Cruz Biotechnology) in a rotary platform for 48 h at 4°C. Membranes were washed thrice with TBST containing 5% (w/v) skimmed milk for 10 min each time. Membranes were incubated in TBST containing 5% (w/v) skimmed milk with mouse IgG kappa binding protein (m-IgGκ BP) conjugated to HRP (1:2000) (sc-516102, Santa Cruz Biotechnology) for 3h at 4°C on a rotary and washed thrice with TBST for 10 min each time. Membranes were developed and imaged as described above. Post imaging, same membranes were probed with anti-beta actin HRP conjugated (sc-47778 HRP, Santa Cruz Biotechnology) as a loading control. Membranes were developed and imaged as above.

### Mouse experiments

National Animal Experiment Board has approved our mouse experiments (salmonellosis - ESAVI/24418/2018, IBD - ESAVI/16359/2019). ***i) Colony breeding***. We used the previously described body-wide Parp14 knockout mice ^20^, kindly provided by Adam Hurlstone (University of Manchester, UK), in a specific-pathogen-free area at Central Animal Laboratory of the University of Turku with free access to soy-free diet and water ad libitum. The Parp14 knockout mice were backcrossed for 10 generations to C57BL/6N background before starting the experiments. Thereafter, heterozygous mice were inbred to generate homozygous littermates for experiments. Genotyping of mice was carried out from DNA extracted from earmarks of two-week-old to three-week-old mice by PCR (primer sequences for wt, 5 -GGCCTAACTATTCACTCGTGT-3 (prAPV426) and 5 -CTGCTCTTCTAGATGATGCAGA-3 (prAPV443); and primer sequences for Parp14 knockout mice, 5 -GATGCAACTGCAAGAGGGTTTAT-3 (LTR rev_comp prAPV437) and 5 -CTGCTCTTCTAGATGATGCAGA-3 (prAPV443). ***ii) DSS exposure.*** Male mice of age 6-8 weeks were used for DSS treatment. They were allocated to two treatment groups, i.e., control and DSS with similar body weight and littermate distribution. DSS (40,000 Da, TdB Consultancy AB, Uppsala, Sweden, #DB001) solution was fresh, dissolved in autoclaved water to a 2.5% (w/v) concentration, and administered to mice in drinking water during days 0-7 of the experiment ^17,18^. Control mice were matched to DSS-treated mice according to age, and starting weight, and were treated equally to DSS-treated mice, except that they did not receive DSS. The mice were weighed daily during the DSS treatment. Stool consistency was also examined, and presence of blood in stool was measured during and after the DSS challenge. The mice were scored for presence of blood in stool (0 = none; 1 = small amounts of blood in stool pellets; 2 = blood found throughout pellet; 3 = clotted blood at anus; 4 = fresh blood on mice or on bedding materials of cage), and stool consistency (1 = normal; 2 = formed but soft; 3 = slightly loose; 4 = liquid). Following treatment (see Fig. 3), mice were sacrificed by CO2 asphyxiation. Blood was collected by cardiac puncture and further processed for flow cytometry. The colon was excised, its length was measured, and then it was washed with PBS and cut into proximal, mid and distal halves, which were studied separately. Spleens were also collected for flow cytometry. Samples for RNA analysis (liquid N2) and histology (4% paraformaldehyde) were collected after colon tissues were cut in half longitudinally. Three variables (body weight, stool consistency, blood in stool) were measured and summarized with descriptive statistics. For analyses, weight was converted to percentages. Differences between the groups were studied by Friedman test and difference of days were studied by Wilcoxon signed rank test for each of the dependent variables (body weight, stool consistency, blood in stool). The normality of variables was evaluated visually and tested with the Shapiro-Wilk test. Due to the non-normality of the continuous variables, nonparametric methods were used. The statistical significance level was set at 0.05 in all tests (two-tailed). The analyses were performed using the SAS system, version 9.4 for Windows (SAS Institute Inc., Cary, NC, USA). ***iii) Salmonella infection.*** *Salmonella* mice experiments were performed according to Barthel and co-workers ^19^. The naturally streptomycin-resistant strain *S. enterica* serovar Tyhphimurium SL1344 was purchased from the Culture Collection University of Gothenburg (CCUG 51871), Gothenburg, Sweden. *Salmonella* bacteria were grown for 12h at 37°C in Luria-Bertani broth with shaking, diluted 1:20 in fresh medium, and sub-cultured for 4h with shaking. Bacteria were washed twice and suspended in ice-cold sterile PBS. Female C57BL/6N mice of age 6-8 weeks were used for *Salmonella* infection experiments. Water and food were withdrawn 4h before per os (p.o.) treatment with 20 mg of streptomycin (75 μl of sterile solution or 75 μl of sterile water [control]). Afterward, animals were supplied with water and food ad libitum. At 20h after streptomycin treatment, water and food were withdrawn again for 4 h before the mice were infected with 10^8^ cfu of *Salmonella* (50 μl suspension in PBS p.o.) or treated with sterile PBS (control). Thereafter, drinking water ad libitum was offered immediately and food 2h post-infection (p.i.). At the indicated times p.i., mice were sacrificed by CO2 asphyxiation, and tissue samples from the intestinal tracts were removed for Parp14 immunohistochemical staining (sample collected in 4% paraformaldehyde) and RNA analysis (sample collected in liquid N2).

### Quantitation of histological changes in the mouse colon tissue

FFPE tissues were cut in longitudinal 5 µm thick sections prior to hematoxylin & eosin (HE) staining. HE staining was performed using standard methods. All samples were scanned using a Pannoramic 1000 Slide scanner (3DHistech, Budapest, Hungary) with 20× objective and analyzed with a Pannoramic viewer (3D Histech, software version 1.15.4). Severity of colonic inflammation was scored according to ^39^.

Briefly, i.) immune cell infiltration: score of 0 = healthy, 1 = some inflammatory cells seen in mucous, 2 = moderate infiltration, and 3 = a severe and acute inflammation; ii) edema scoring was based on location and severity of edema and ranged from 0–3; iii) erosion depth: 0 = no erosion, 1 = erosion of epithelium, 2 = erosion of mucous, 3 = erosion going through muscular lamina; iv) loss of goblet cells: 0 = no change, 1 = weak, 2 = moderate, and 3 = severe effect/lot. Two people performed the scoring independently and blind for the sample group. Presented scores are averages of these two analyses. Variables edema, erosion, goblet cell loss and immune infiltration were measured for distal and proximal colon. Variable colon length was measured in millimeter (mm) and mean of measurements from each group were calculated. Variables were summarized with descriptive statistics and associations between group and variables were studied by Kruskal-Wallis test and Dwass-Steel-Chritchlow-Fligner test for post hoc comparisons. The normality of variables was evaluated visually and tested with the Shapiro-Wilk test. Due to the non-normality of the continuous variables, nonparametric methods were used. The statistical significance level was set at 0.05 in all tests (two-tailed). The analyses were performed using the SAS system, version 9.4 for Windows (SAS Institute Inc., Cary, NC, USA).

### Mouse Parp14 immunohistochemistry

***i) Anti-Parp14 IHC.*** Mice (C57BL/6N and FVB/n) FFPE tissue sections were stained with anti-Parp14 antibody in the similar way as done for human FFPE biopsy specimens as mentioned above. ***ii) Double immunofluorescence*.** Anti-F4/80 (rat anti-mouse MCA497A647, Biorad, dilution 1:500) and anti-Parp14 (sc377150, Santa Cruz Biotechnology, dilution 1:500) were mixed in 5% (w/v) BSA in TBST and added to colon sections post blocking for overnight incubation at 4°C in a humidified chamber. Anti-F4/80 secondary antibody was Alexa-fluor 647 conjugated. Secondary antibody Alexa-fluor 488 goat anti-mouse IgG (H+L) (A11001, Invitrogen, dilution 1:1000) was used to visualize mouse anti-Parp14. Tissue sections were washed with PBST and counterstained using 300 nM of DAPI (sc-3598, Santa Cruz) for 1min. Mounting was done using Histo-Clear and sections were kept in dark at 4°C before imaging using Zeiss AxioImager M1. DAPI (nucleus), GFP (green, Parp14) and Alexa-fluor 660 (red, F4/80) channels were used in image acquisition.

### Isolation of RNA and quantitative PCR analysis

Total RNA was isolated from mouse tissues using TRIsure reagent (BIO-38033, Bioline Gmbh, Germany) and genomic DNA was digested using RNase-free DNase (rDNase) from Machery-Nagel as per the manufacturer’s instructions. Briefly, colon tissue was homogenized using stainless steel balls (IKA 5mm stainless steel balls, Fisher scientific) and compact bead mill (TissueLyser LT, Qiagen). Colon tissue was placed in TRIsure reagent during homogenization. After this, chloroform was used for phase separation. RNA in the upper aqueous phase was precipitated using isopropyl alcohol. Pellet was washed with 70% ethanol and dissolved in nuclease free water. Dissolved RNA was mixed with rDNase and rDNase reaction buffer as per manufacturer’s instructions. This mixture was incubated at 37°C for 10 min. Post-gDNA digestion, RNA was precipitated using 3M sodium acetate, pH 5.2 and 96% ethanol. Pellet was washed with 70% ethanol and dissolved in RNase free water. RNA purity and concentration were measured using DeNovix DS-11 spectrophotometer (Wilmington, DE, USA). 1µg of dissolved RNA was reverse transcribed using SuperScript III reverse transcriptase (#1808044, ThermoFisher Scientific) and Oligo (dT) 12-18 Primer (#18418012, ThermoFisher Scientific). Separate real-time PCR (RT-PCR) was carried out in duplicates using TaqMan gene expression assays (Applied Bio systems, Foster city, Ca) for *parp14* (Assay ID: Mm00520984_m1) and the reference gene, glyceraldehyde-3-phosphate dehydrogenase, *gapdh* (Assay ID: Mm99999915_g1) on the Rotor-Gene Q real-time PCR cycler (Qiagen). Thermal cycling conditions included an initial denaturation step at 95°C for 10 min followed by 40 cycles of 95°C for 15s and 60°C for 1 min. Relative mRNA levels were determined using the 2^-ΔΔCT^ method with GAPDH as reference. If the standard deviation of duplicate Ct values from each sample was 0.5 or more, those samples were not used for statistical analysis. Statistical analysis of 2^-ΔΔCT^ values were calculated using unpaired t test (two tailed).

### Bacterial community analysis of mouse fecal sample

DNA of fecal pellets was extracted using Machery-Nagel NucleoSpin DNA Stool kit (reference no. 740472.250) using the manufacturer’s protocol. The 16S rRNA gene amplicons were sequenced at University of Illinois, Roy J. Carver Biotechnology Center, DNA Services Laboratory. In brief, the 16S rRNA gene amplicons were generated with the barcoded Full-Length 16S rRNA gene primers (forward: AGRGTTYGATYMTGGCTCAG, reverse: RGYTACCTTGTTACGACTT) from PacBio and the 2x Roche KAPA HiFi Hot Start Ready Mix. Amplicons were converted to a library with the SMRTBell Express Template Prep kit 2.0. The pooled library was sequenced on 3 SMRTcell 8M on a PacBio Sequel IIe using the CCS sequencing mode and a 12hs movie time. The raw sequence data has been deposited in at NCBI under the Bio project accession, PRJNA977978. A total of three DNA extractions and one PCR water controls were also sequenced along with the samples. The bioinformatics analysis was performed using mothur (v1.48.0) software. The raw 16S rRNA data were processed as described in ^40^ with modifications needed for the PacBio sequence analysis. In brief, fastq.info command in which PacBio parameter set to be true, was used to convert the fastq files to fasta files. The command of make.group was used to assign each sequence to its sample. Subsequently, merge.files command was used to merge fasta files. The screen.seqs was used to remove the ambiguity and homopolymer (maximum eight) sequences. The sequences were aligned using Silva reference database (v138) followed by removal of potential chimeras. Subsequently, sequences were clustered at 97% similarity level to create an operational taxonomic units (OTU) abundance table. The representative sequences of each OTUs were classified using a mothur compatible Ribosomal Database Project (v18). All the statistical analysis were performed using R software (v4.1.1; R Core Team, 2020). A total of 954 OTUs were obtained across the five wt and five knock-out mutant mouse samples. Two blank DNA isolation controls produced 47 OTUs. The number of sequences in each sample ranged from 2058 to 24673. The subsampling for the equal sequencing depth (2058 sequences per sample) resulted in 524 OTUs. Among the 47 OTUs in blank DNA isolation controls, only one OTU (Otu0063) was found in the fecal pellet samples. The subsampling resulted in removal of the potential contaminant Otu0063 from the fecal pellet samples.

### Flow cytometry-based leukocyte profiling

***i) Cell isolation.*** The mid-colon was excised into 10 mL of ice-cold PBS and surrounding fat tissue removed. The colon was cut open longitudinally and rinsed in PBS to remove fecal content. Colon was further cut into 5-10 mm pieces and put into a 15 mL falcon tube with 5 mL prewarmed 2 mM EDTA in Hank’s balanced salt solution (HBSS, Gibco) and shaken at 250 rpm for 15 min at 37°C. Tube was manually shaken vigorously and supernatant removed while tissue segments recovered with colander. 5 mL of warm 2 mM EDTA in HBSS was added on the segments, incubated for 30 min at 37 °C with shaking 250 rpm, and then shaken vigorously by hand. The epithelial cell fractions were discarded, and the lamina propria tissues were digested in 5 mL of enzyme cocktail containing 1 mg/mL Collagenase VIII (Sigma C2139) and 10 μg/mL Dnase I (D5025) diluted in RPMI (Corning) with 1 % Fetal Calf Serum (FCS, BioWest, S181B) at 250 rpm for 45 min at 37°C. Digestive enzyme activity was stopped by adding 5 mL cold FACS buffer (PBS, 2 % FCS, 1 mM EDTA). Digested tissue was then filtered through 70 μm cell strainer, centrifuged at 1500 rpm for 5 min, and stained for flow cytometry analysis. From the blood sample, 100 µl was taken to a tube containing a drop (approximately 10 µl) of heparin (Heparin LEO 100 IU/mL, LEO Pharma). Samples were kept on ice and erythrocytes lysed with hypotonic treatment. First 1 mL of 0.2 % NaCl was added per sample, vortexed up to 15 seconds and then 1 mL of 1.6 % NaCl was added. Cells were pelleted (1006 x g, 1.5 minutes) and lysis repeated in total of four times. Finally, cells were washed twice with 1 mL of PBS and filtered through silk with 77 µm pore size. Spleens were collected to PBS on ice. Spleens were minced through a metal mesh using the plunger of 1 mL syringe. Mesh was washed with 1 mL PBS and the cell suspension was collected and pelleted (1006g, 1.5 minutes). To remove red blood cells from suspension, cells were lysed with a NaCl-treatment by first adding 0.2 % NaCl and vortexing 15 seconds and then adding 1.6 % NaCl. Cell suspension was pelleted (1006g, 1.5 minutes) and washed with PBS. Finally, cells were eluted to PBS and filtered through silk with 77 µm pore size. ***ii) Flow cytometry.*** Cells were stained with Fixable Viability Dye eFluor 780 (eBioscience, 65-0865) to label the dead cells. Unspecific binding to low-affinity Fc-receptors was blocked by incubating the cells with unconjugated CD16/32 antibody (BioXCell, clone 2.4G2). Cells were subsequently stained for 30 min at 4 °C with antibodies diluted in the FACS buffer. Immune cell panel included CD45-PerCP-Cy5-5 (Becton Dickinson (BD), #550994), CD4-PE (BD, #553049), CD8-Bv650 (BD, #563234), B220-PE-CF594 (BD, #562313), CD11b-BB515 (BD, #564454), CD11c-BV711 (BioLegend, #117349), Ly6c-BV421 (BD, #562727), Ly6G-BV510 (BioLegend, #127633), SiglegF-A647 (BD, #562680), CD64-BV786 (BD, #741024) and F4/80-PE-Cy7 (Invitrogen, #25-4801-82). Cells were washed with FACS buffer and fixed with 1% formaldehyde in PBS. Samples were acquired with LSRFortessa flow cytometer with FACSDiVaTM version 8 software (Becton Dickinson), and data were analyzed with the FlowJo software (FlowJo LLC). The gating strategies for flow cytometric analyses are depicted in Fig. S7A. Results were statistically analyzed in GraphPad Prism.

### Mouse colon tissue bulk RNA-Seq and data analysis

Total RNA was extracted as described above. Preparation of RNA library and transcriptome sequencing was conducted by Novogene Co., LTD (Cambridge, UK). ***i) RNA quality.*** RNA integrity was assessed using the RNA Nano 6000 Assay Kit of the Bioanalyzer 2100 system (Agilent Technologies, CA, USA). ***ii) Library preparation for transcriptome sequencing.*** Briefly, mRNA was purified from total RNA using poly-T oligo-attached magnetic beads. Fragmentation was carried out using divalent cations under elevated temperature in First Strand Synthesis Reaction Buffer(5X). First strand cDNA was synthesized using random hexamer primer and M-MuLV Reverse Transcriptase (RNase H-). Second strand cDNA synthesis was subsequently performed using DNA Polymerase I and RNase H. Remaining overhangs were converted into blunt ends via exonuclease/polymerase activities. After adenylation of 3’ ends of DNA fragments, Adaptor with hairpin loop structure were ligated to prepare for hybridization. To select cDNA fragments of preferentially 370-420 bp in length, the library fragments were purified with AMPure XP system (Beckman Coulter, Beverly, USA). Then PCR was performed with Phusion High-Fidelity DNA polymerase, Universal PCR primers and Index (X) Primer. At last, PCR products were purified (AMPure XP system) and library quality was assessed on the Agilent Bioanalyzer 2100 system. ***iii) Clustering and sequencing.*** The clustering of the index-coded samples was performed on a cBot Cluster Generation System using TruSeq PE Cluster Kit v3-cBot-HS (Illumia) according to the manufacturer’s instructions. After cluster generation, the library preparations were sequenced on an Illumina Novaseq platform and 150 bp paired-end reads were generated. ***iv) Data analysis - Quality control.*** Raw data (raw reads) of fastq format were firstly processed through Novogene’s in-house perl scripts. In this step, clean data (clean reads) were obtained by removing reads containing adapter, reads containing ploy-N and low quality reads from raw data. At the same time, Q20, Q30 and GC content the clean data were calculated. All the downstream analyses were based on the clean data with high quality. ***v) Data analysis - Reads mapping to the reference genome.*** Reference genome and gene model annotation files were downloaded from genome website directly. Index of the reference genome was built using Hisat2 v2.0.5 and paired-end clean reads were aligned to the reference genome using Hisat2 v2.0.5. We selected Hisat2 as the mapping tool for that Hisat2 can generate a database of splice junctions based on the gene model annotation file and thus a better mapping result than other non-splice mapping tools. ***vi) Data analysis - Quantification of gene expression level.*** The featureCounts v1.5.0-p3 was used to count the reads numbers mapped to each gene. Subsequently, FPKM of each gene was calculated based on the length of the gene and reads count mapped to this gene. FPKM, expected number of Fragments Per Kilobase of transcript sequence per Millions base pairs sequenced, considers the effect of sequencing depth and gene length for the reads count at the same time, and is currently the most commonly used method for estimating gene expression levels. ***vii) Data analysis - Differential expression analysis.*** Differential expression analysis of two conditions/groups (two biological replicates per condition) was performed using the DESeq2 R package (1.20.0). DESeq2 provide statistical routines for determining differential expression in digital gene expression data using a model based on the negative binomial distribution. The resulting *p*-values were adjusted using the Benjamini and Hochberg’s approach for controlling the false discovery rate. Genes with an adjusted *p*-value <0.05 found by DESeq2 were assigned as differentially expressed. ***viii) Data analysis - GO enrichment analysis of differentially expressed genes (DEGs).*** This part of the data analysis was performed in-house. To analyze cellular and physiological associations of the up-and down regulated DEGs, we performed a Gene Ontology (GO) enrichment analysis (https://geneontology.org/docs/go-enrichment-analysis/). Only those DEGs that passed our in-house filtering criteria were used in the GO enrichment analysis, i.e., up-regulated genes, log2(FoldChange) > 1 & padj < 0.05; down-regulated genes, log2(FoldChange) < -1 & padj < 0.05. The GO biological process and cellular component annotation data set was used with the binomial test type using Bonferroni correction for multiple testing. The raw RNA-Seq data has been deposited to Gene Expression Omnibus public functional genomics data repository (https://www.ncbi.nlm.nih.gov/geo/) with accession number GSE252812.

### Human bulk colon tissue transcriptome analysis

Bulk transcriptome data GEOD-14580^24^ and E-GEOD-4183^25^ was accessed using the ArrayExpress repository (https://www.ebi.ac.uk/biostudies/arrayexpress) and biogps.org gene annotation portal ^41^. Using biogps.org, Parp14 mRNA expression value (fluorescence intensity) for each patient from the two datasets was downloaded. Using sample information from ArrayExpress, sample groups were assigned and mean of expression value was calculated two draw separate graphs. The difference between two groups was statistically measured using Mann-Whitney test.

## SUPPLEMENTARY FIGURE LEGENDS

**Figure S1.**
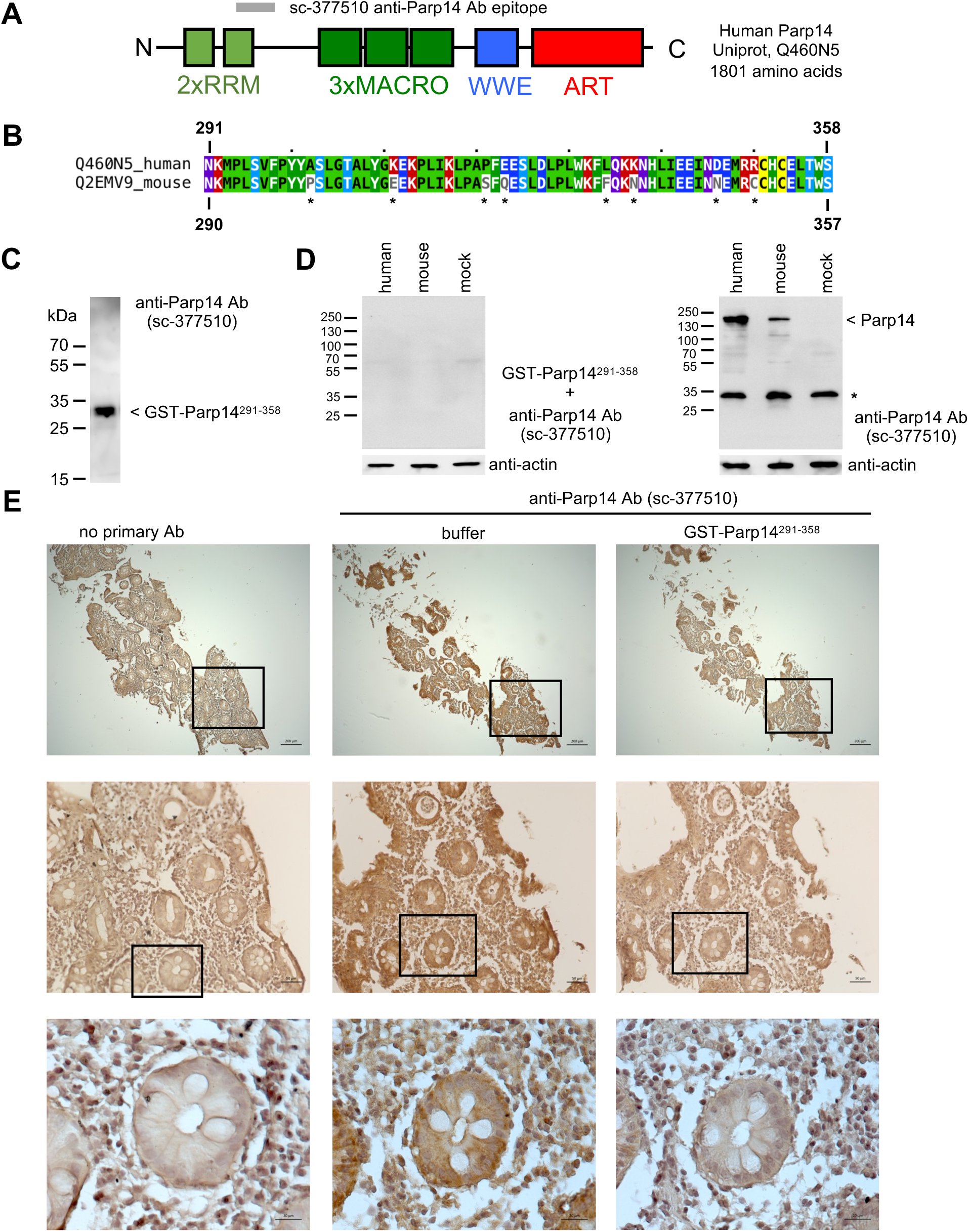
Specificity of the anti-Parp14 antibody. **A)** The domain structure of Parp14. The localization of the human epitope used to obtain the mouse monoclonal anti-Parp14 antibody (sc-377150, Santa Cruz Biotechnology) is shown with a gray bar. RRM, RNA recognition motif domain; MACRO, macrodomain; WWE, tryptophan-tryptophan-glutamate domain; ART, ADP-ribosyltransferase domain. **B)** Similarities of human and mouse Parp14. This area refers to the epitope of human Parp14 that was used to obtain the mouse monoclonal anti-Parp14 antibody. **C)** Western analysis of the purified GST-Parp14^291–358^ (human). Expected size of the fusion protein is 34.85 kDa. **D)** Epitope competition Western analysis. Full length human and mouse Parp14 were ectopically expressed in HEK293T cells. The cell lysates were analyzed for Parp14 expression in the presence or absence of GST-Parp14^291–358^. The star refers to a protein, which consistently stained positive in anti-Parp14 Western analysis of HEK293T cells (see also Fig. S3). **E)** Epitope competition IHC analysis. IHC staining of Parp14 in an FFPE endoscopic biopsy section of an UC patient (TCML_VM_28A, ascending colon, see Suppl. data file 1). 5x, 20x and 40x oil objective images are shown (1:10000 dilution of anti-Parp14 antibody).

**Figure S2.**
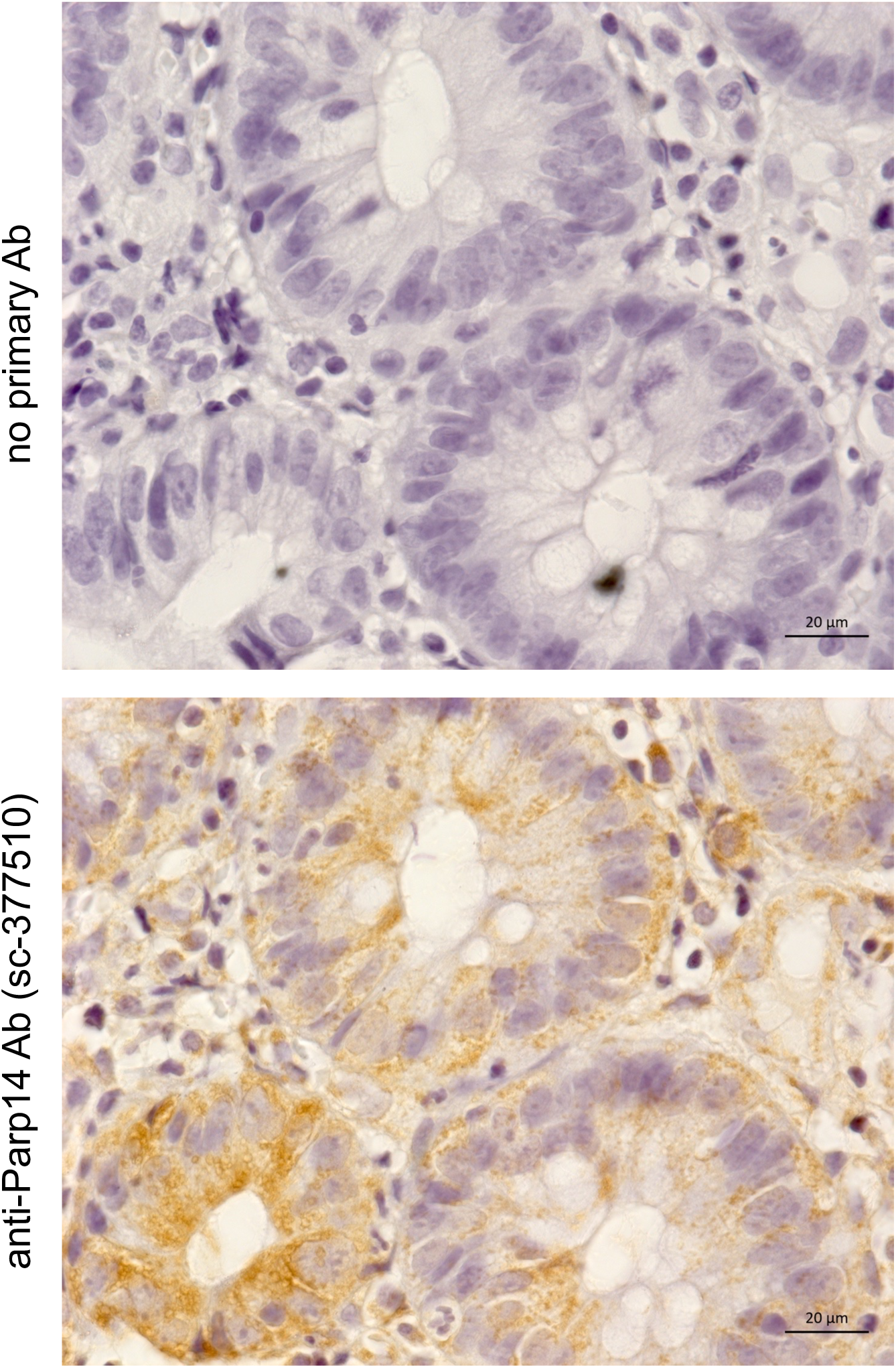
Granular staining pattern of Parp14 in the epithelial cells of human colon. IHC staining of Parp14 in an FFPE endoscopic biopsy section of an UC patient (TCML_VM_21A, ascending colon, see Suppl. data file 1). 63x objective images are shown (1:10000 dilution of anti-Parp14 antibody).

**Figure S3.**
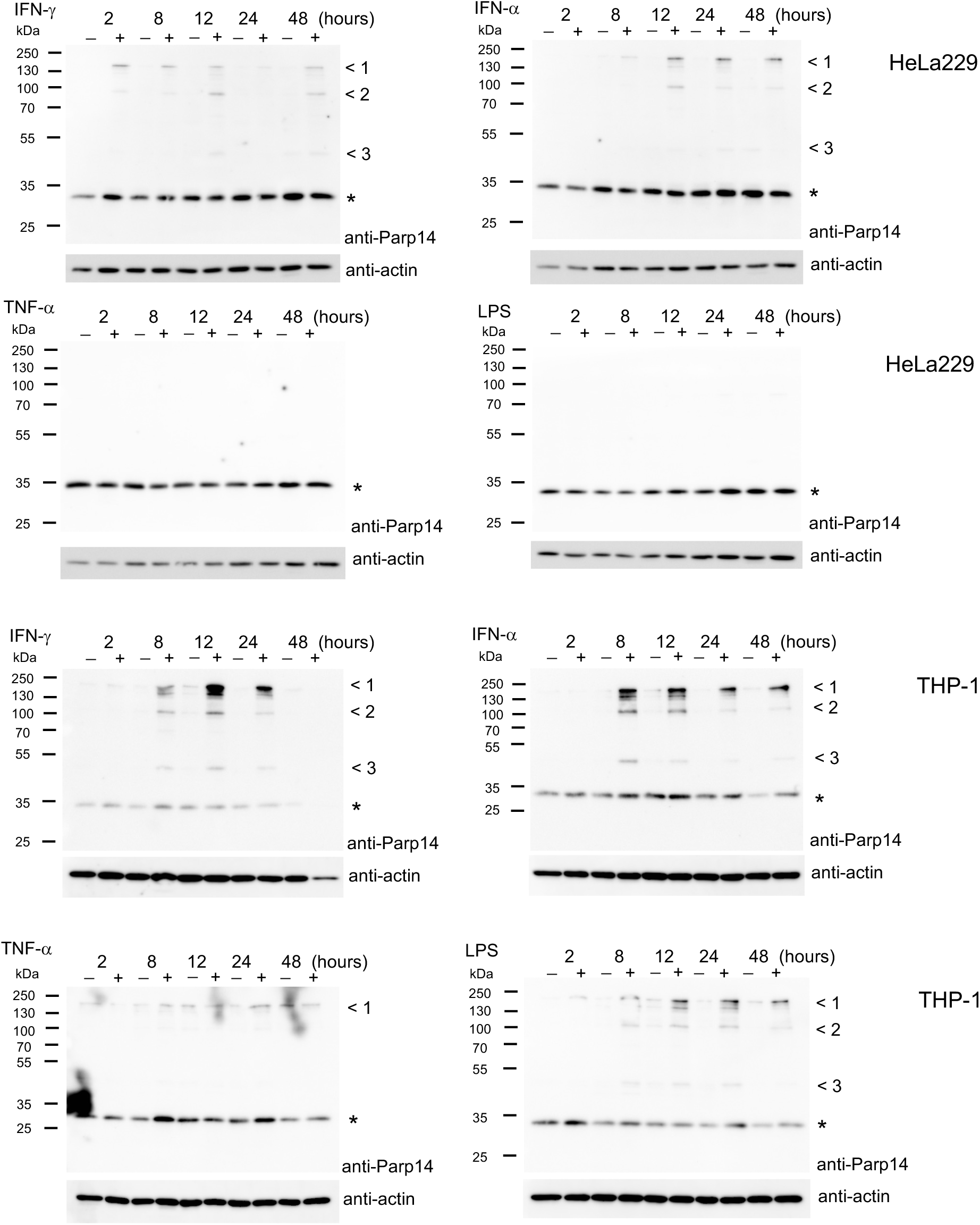
Expression of Parp14 in cultured epithelial cells and macrophages. The cells were either treated with different inflammatory stimuli or left untreated for 2-48 hours (100 U/mL of IFN-γ, 100 U/mL of INF-α, 10 ng/mL of TNF-α or 100 ng/mL of LPS). The cell lysates were analyzed for Parp14 expression using the mouse monoclonal anti-Parp14 antibody. Data with maximum exposure time of 20 min is shown. Loading was controlled with anti-actin Western analysis. Expected size of human Parp14 is 202.83 kDa (Uniprot Q460N5-6, canonical isoform). Arrow 1, full length Parp14; Arrows 2-3, truncated forms or isoforms of Parp14. The star refers to a protein, which consistently stained positive in anti-Parp14 Western analysis with both cell lines.

**Figure S4.**
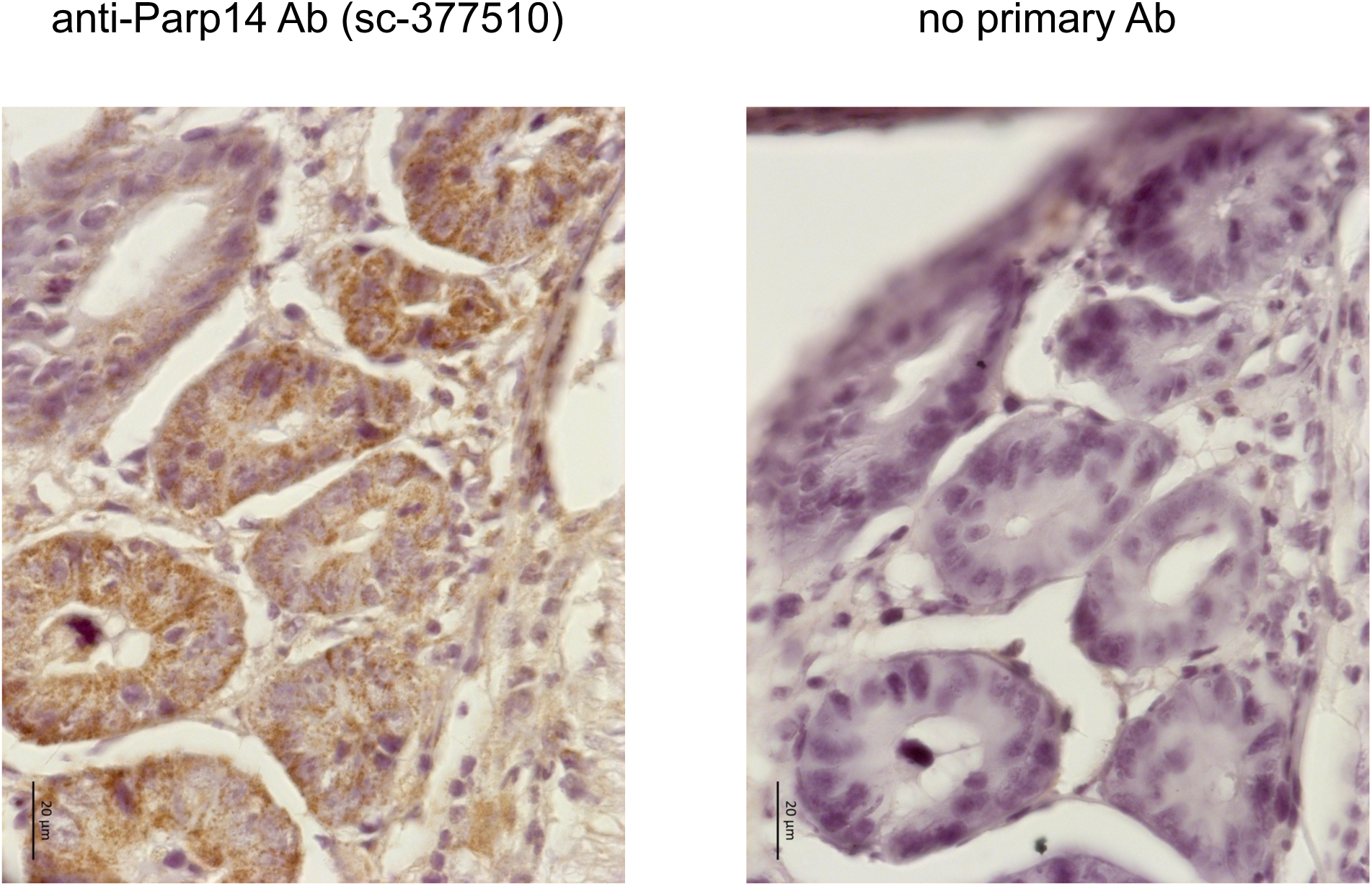
Granular staining pattern of Parp14 in the epithelial cells of mouse colon. IHC staining of Parp14 in an FFPE tissue section of a DSS-treated (2.5%, 7 days) FVB/n mice. 63x objective images are shown (1:500 dilution of anti-Parp14 antibody).

**Figure S5.**
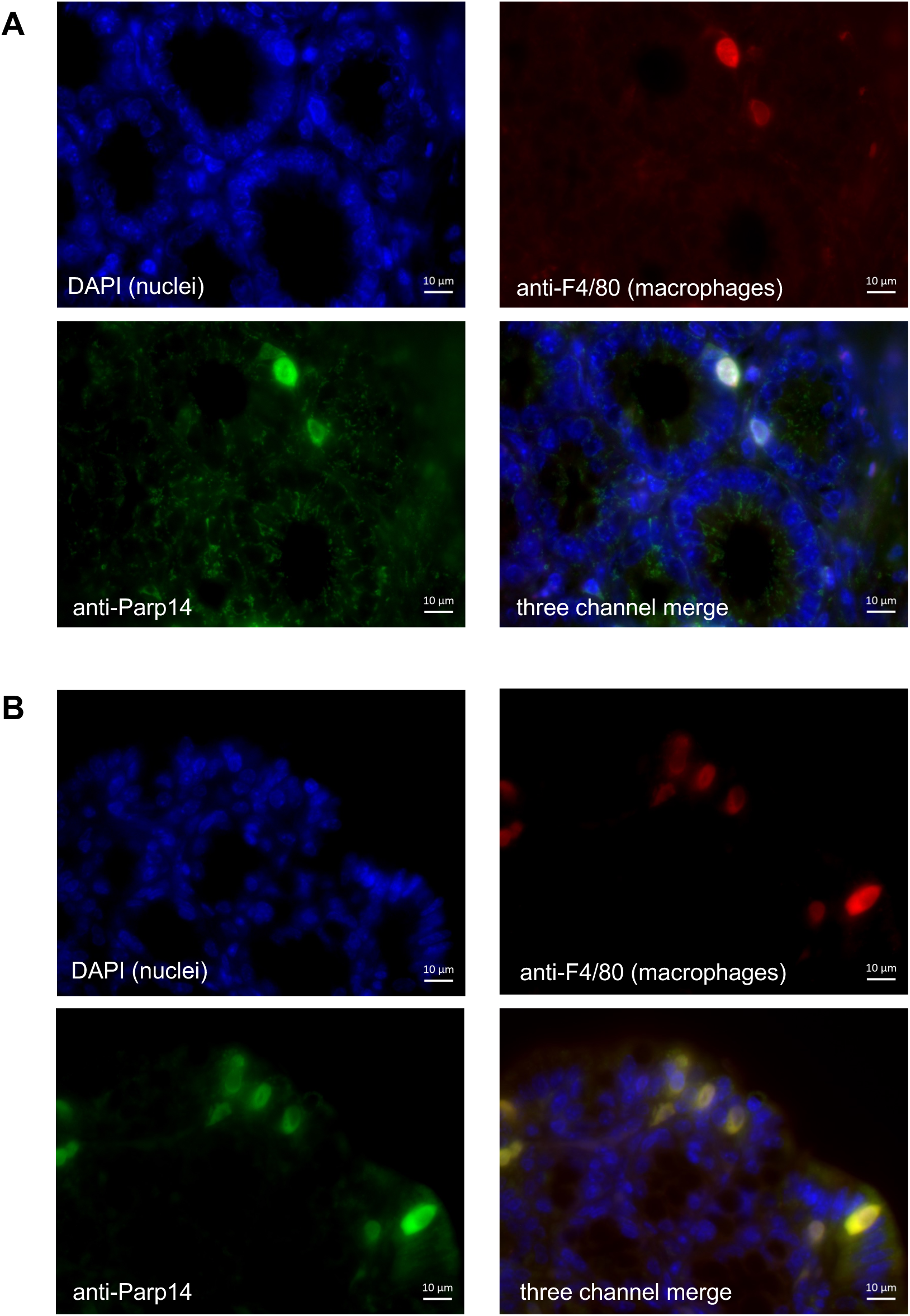
Immunofluorescence imaging of macrophage marker F4/80 and Parp14 in the mouse colon. Two different FFPE colon sections (**A-B**) derived from the DSS treated C57BL/6N mice (2.5%, 7 days) were analyzed for Parp14 (green channel) and macrophage marker F4/80 (red channel). 63x objective images are shown (1:500 dilution of anti-Parp14 antibody).

**Figure S6.**
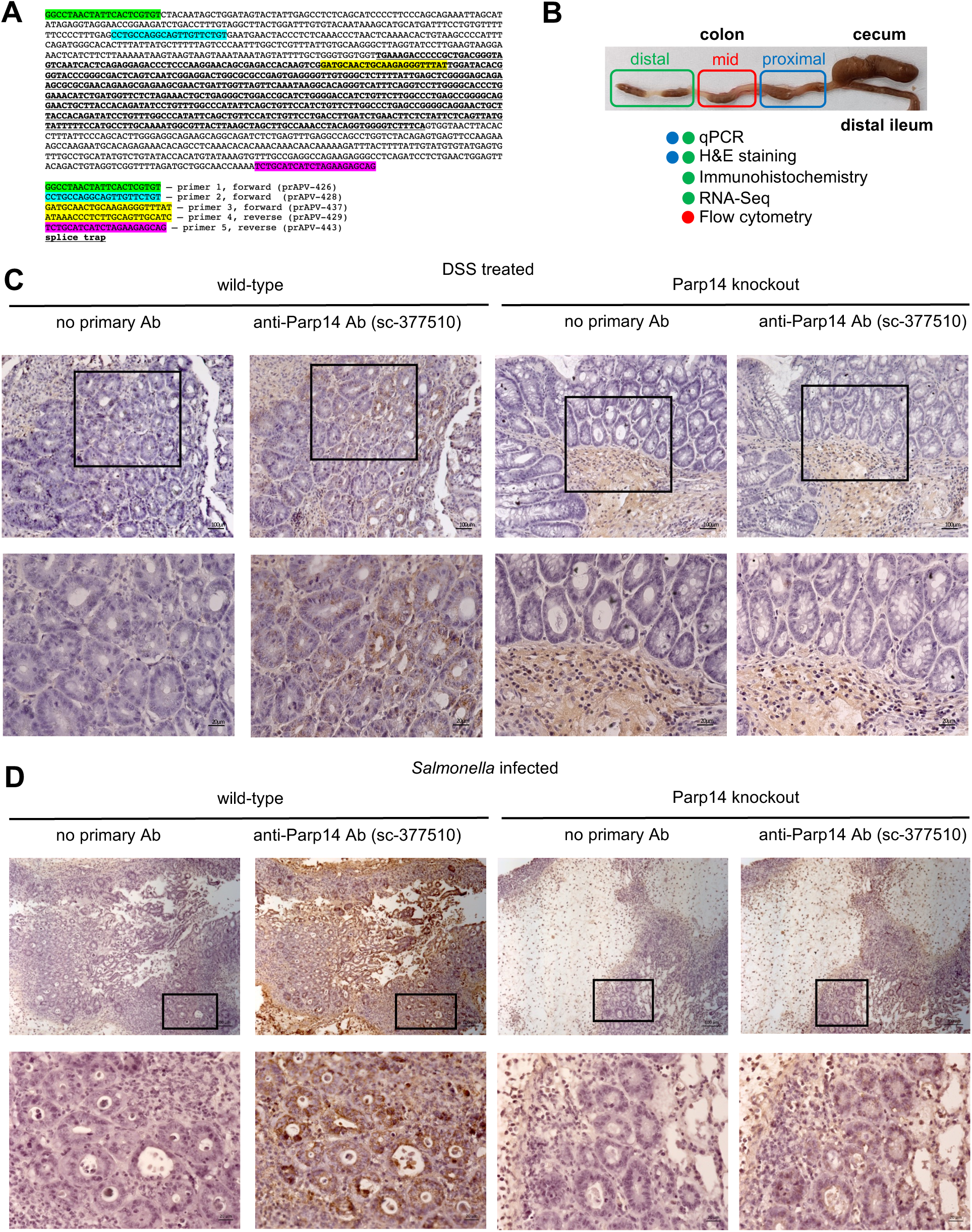
Parp14 expression and localization in the mouse colon in two intestinal inflammation models – verification of Parp14 deficiency in the Parp14 knockout mice. **A)** Nucleotide sequence of the location of the gene trap-targeted mouse *parp14* allele. The gene trap is located between exons 1 and 2. Sequences of the oligonucleotide primers used to genotype the mice during breeding and to sequence the gene trap-targeted allele are shown. **B)** Schematic representation of the sampling for different downstream analysis. **C**) DSS colitis IHC data. Parp14 IHC of distal colon of DSS (2.5%, 7 days) treated wt and Parp14-deficient C57BL/6N mice. 20x and 40x objective images are shown (1:500 dilution of anti-Parp14 antibody). **D**) Salmonellosis IHC data. Parp14 IHC of proximal colon of *Salmonella* infected (5 days) wt and Parp14-deficient C57BL/6N mice. 10x and 40x objective images are shown (1:500 dilution of anti-Parp14 antibody).

**Figure S7.**
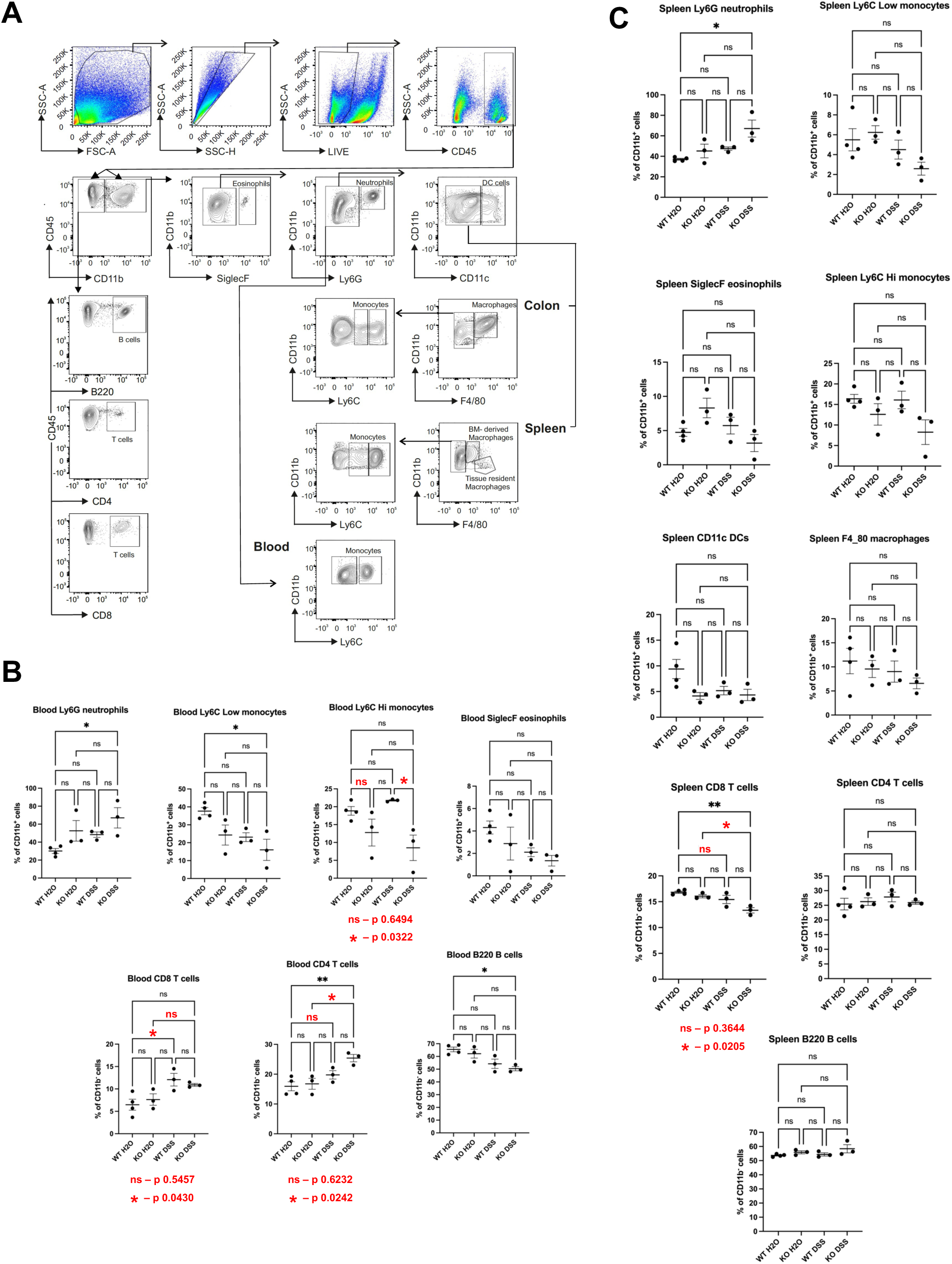
DSS-treated Parp14-deficient mice have abnormal numbers of monocytes and T cells in their blood and spleen. Blood and spleen single-cell suspensions post-DSS and control treatment at day 8 were stained for myeloid-derived cells, i.e., granulocytes (neutrophils and eosinophils), monocytes, macrophages, dendritic cells, and lymphoid-derived cells (B and T cells). **A)** Gating strategy, and results with **B)** blood and **C)** spleen. Each dot represents one mouse. Data are presented as mean ±SEM. Statistical analysis is done with one-way ANOVA with the Bonferroni post-hoc test.

**Figure S8.**
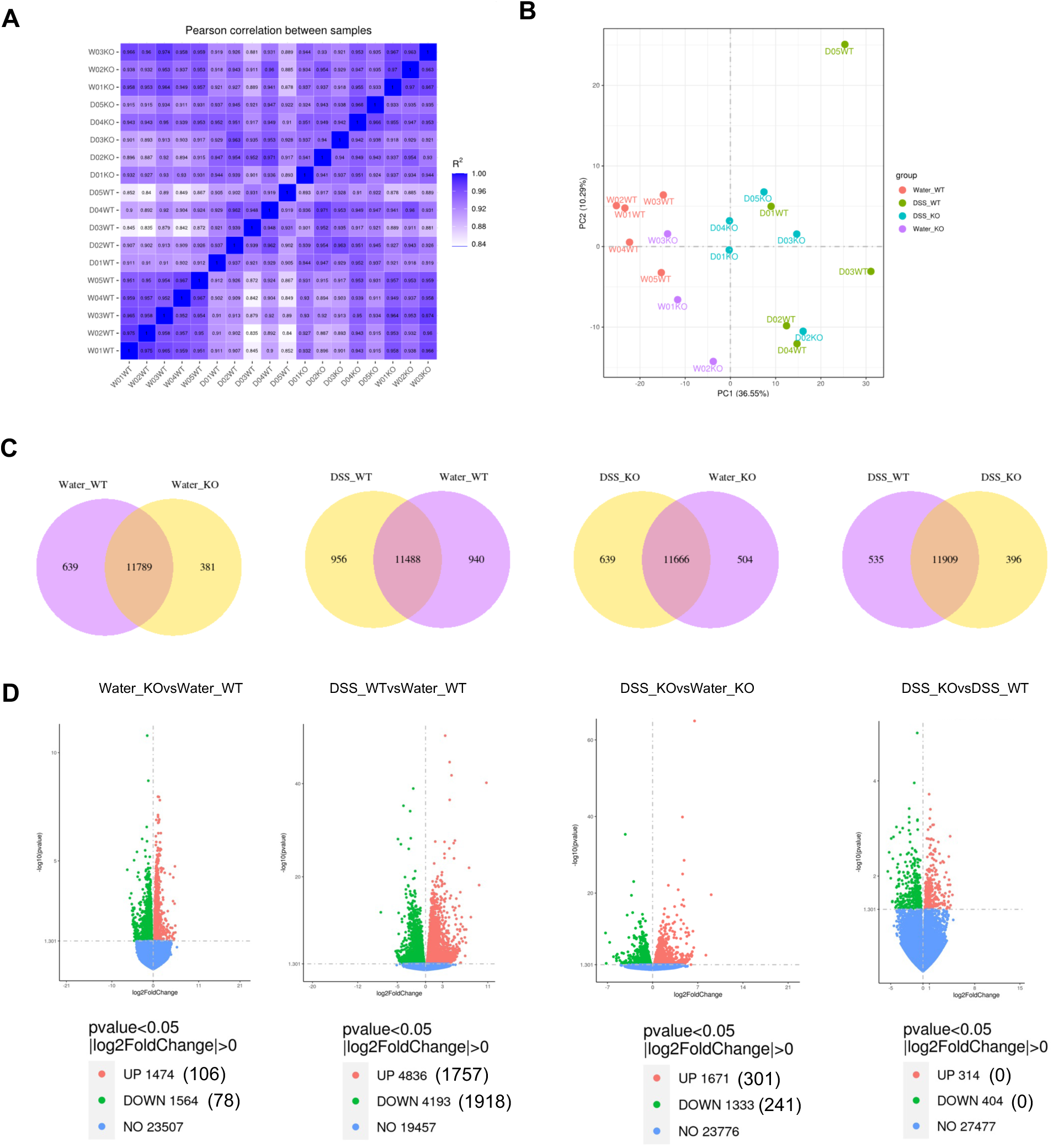
The mid-colon bilk tissue RNA-Seq analysis. Data reliability metrics on the different biological replicates (**A-B**). Numbers of genes detected to be expressed in the four compare groups, as well as their differential gene expression metrics (**C-D**). **A)** Inter-sample correlation heatmap based on FPKM (Fragments Per Kilobase of transcript per Million mapped reads) gene expression values. R^2^ is the square of Pearson correlation coefficient (R). R^2^ values reflect relatively well the different treatments, e.g., the R^2^ values within the water treated wt group are all >0.95 whereas frequently <0.9 when the water treated wt group is compared to the DSS treated wt group. Also, the R^2^ values are always >0.9 within the samples of the four compare groups. **B)** Principal component analysis (PCA) based on FPKM gene expression values. Water- and DSS-treated groups are separated whereas some overlap is visible between the genotypes within both of the treatments. The clear outlier (D05WT) was included into the downstream analysis because of its R^2^ values were always >0.9 with the rest of the water treated wt mice (see Fig. S8A). **C)** The Venn diagram of shared and unique genes that were detected to be expressed among the four compare groups. The integer is the number of shared genes. **D)** Volcano plots of differentially expressed genes (DEGs). Specific information of the DEGs is given in Suppl. data file 2. The x-axis shows the fold difference in gene expression between different samples, and the y-axis shows the statistical significance of the differences. Red dots represent up-regulation genes and green dots represent down-regulation genes. The dashed line indicates the threshold line for statistically significant differential gene expression. The values in brackets refer to the number of DEGs that were used for all the subsequent data analysis after our in-house filtering, i.e., UP genes, log2(FoldChange) > 1 & padj < 0.05; DOWN genes, log2(FoldChange) < -1 & padj < 0.05 (Suppl. data file 2).

**Figure S9.**
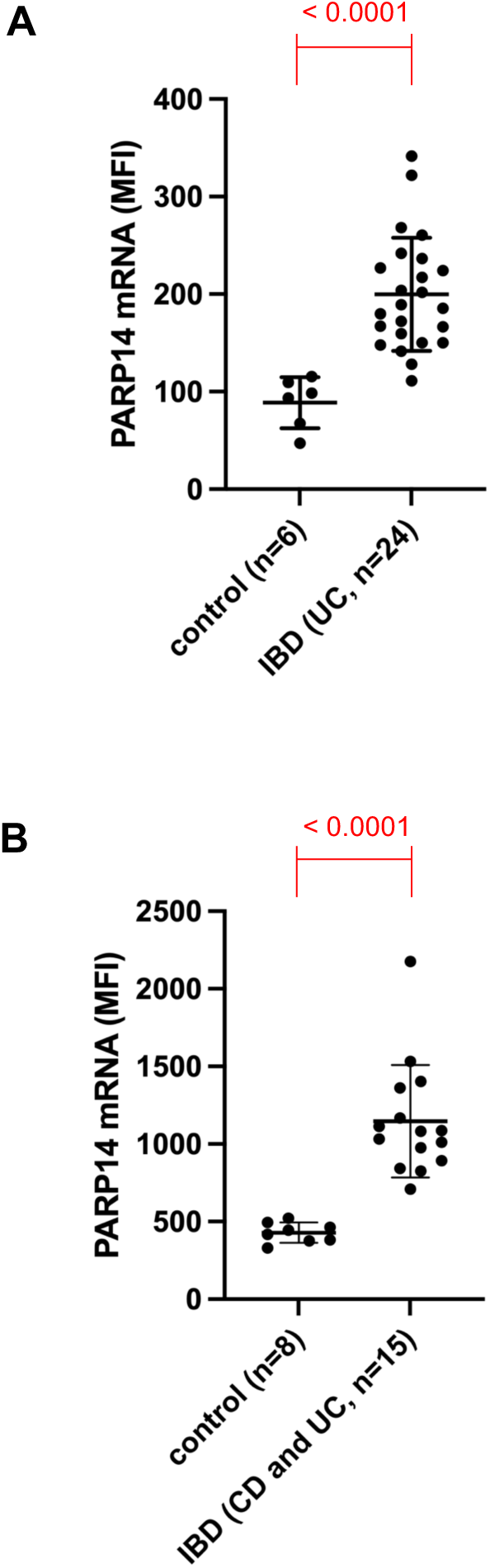
Elevated amount of Parp14 transcripts in IBD patients. Publicly available transcriptome data of colon biopsies from two IBD cohorts are shown for Parp14 transcript levels, i.e., **A)** E-GEOD-14580, and **B)** E-GEOD-4183. Each dot represents a single individual (means with standard deviation, statistics with Mann-Whitney test).

